# Widespread cryptic variation in genetic architecture between the sexes

**DOI:** 10.1101/2021.02.20.432102

**Authors:** Wouter van der Bijl, Judith E. Mank

**Affiliations:** Department of Zoology and Biodiversity Research Centre, University of British Columbia, Canada; Department of Genetics, Evolution and Environment, University College London, UK

**Keywords:** sexual dimorphism, genetic architecture, between-sex genetic correlation, rFM, knock-out

## Abstract

The majority of the genome is shared between the sexes, and it is expected that the genetic architecture of most traits is shared as well. This common architecture has been viewed as a major source of constraint on the evolution of sexual dimorphism (SD). SD is nonetheless common in nature, leading to assumptions that it results from differential regulation of shared genetic architecture. Here, we study the effect of thousands of gene knock-out mutations on 202 mouse phenotypes to explore how regulatory variation affects SD. We show that many traits are dimorphic to some extent, and that a surprising proportion of knock-outs have sex-specific phenotypic effects. Many traits, regardless whether they are monomorphic or dimorphic, harbor cryptic differences in genetic architecture between the sexes, resulting in sexually discordant phenotypic effects from sexually concordant regulatory changes. This provides an alternative route to dimorphism through sex-specific genetic architecture, rather than differential regulation of shared architecture.

## Introduction

In organisms with separate sexes, different evolutionary interests of males and females can lead to divergent trait optima, which can be realized through the evolution of sexual dimorphism. For traits to change from monomorphic to dimorphic, the underlying genetic mechanisms need to be decoupled between males and females. However, even in species with sex chromosomes, males and females share the vast majority of their genome (Bachtrog *et al.*, 2014), leading to the expectation that traits are controlled by the same loci in both sexes (Lande, 1980). This shared genomic architecture is typically considered a source of significant constraint on the evolution of dimorphism (Stewart & Rice, 2018), as traits would need to first become genetically decoupled between females and males before divergence can occur (Lande, 1980; Poissant *et al.*, 2010; Hermansen *et al.*, 2018). Shared trait architecture can lead to intra-locus sexual conflict (Rice & Chippindale, 2001), where alleles at a locus have different fitness effects in males and females, and is this assumed to limit the degree to which the sexes can achieve their respective fitness optima (Hansen, 2006). Indeed, the constraints on the evolution of sexual dimorphism (SD) are often considered both pervasive and persistent, resulting in enduring sexually antagonistic selection on many traits (Rice & Chippindale, 2001; Chenoweth *et al.*, 2008; Poissant *et al.*, 2010; Ruzicka *et al.*, 2019). This persistent constraint is however difficult to reconcile with the fact that sexual dimorphism evolves rapidly (Stewart & Rice, 2018), is seen in a broad array of traits, and differs markedly among related species (Owens & Hartley, 1998).

It has been suggested that sexual dimorphism arises from regulatory differences between males and females (Ellegren & Parsch, 2007; Mank, 2017), and there are good examples of this (e.g. Galouzis & Prud’homme, 2021). Indeed, recent genome-wide scans in fruit flies have shown that protein coding sequence differences are overrepresented among evolutionarily persistent variants thought to be maintained by sexual antagonism (Ruzicka *et al.*, 2019). This might suggest that conflict over coding sequence variation is much harder to resolve compared to conflict over gene expression. However, functional studies have revealed that the genes underlying some dimorphisms are not expressed differently between the sexes (Khila *et al.*, 2012). This indicates that sex-biased expression alone cannot explain all dimorphism, and other mechanisms may exist.

Another perspective on the genetics of sexually dimorphic traits stems from investigations grounded in quantitative genetic theory (Lande, 1980). By comparing the phenotypes of individuals of known relatedness, usually through breeding designs or pedigrees, one can estimate the between-sex genetic correlation (*r_fm_*) for a trait of interest. This correlation describes the extent to which a particular genotype affects both male and female phenotypes in the same way. If *r_fm_* ≈ 1, genotypes affect males and females similarly (i.e. brothers and sisters look alike), while if *r_fm_* ≈ 0, male and female phenotypes vary independently (Lande, 1980). This estimate of *r_fm_* is based on autosomal additive standing genetic variation and measures the additive effects of the many genetic variants that exist in that population at that time. It can therefore be used to predict the extent to which a population can respond to sexually divergent selection. Since this *r_fm_* estimate is based on the additive genetic variance, we will denote it here as 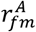 for clarity.

Average estimates of 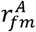 are often close to one (Poissant *et al.*, 2010), suggesting that there is little standing sex-specific genetic variation. However, these estimates are also interpreted by many to reflect the extent to which the autosomal genetic architecture underlying the trait is shared between the sexes (Chenoweth *et al.*, 2008; Poissant *et al.*, 2010; Griffin *et al.*, 2013; e.g. Stewart & Rice, 2018). In other words, a strongly positive 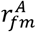 is interpreted to mean that the gene network that produces the phenotypic trait value is largely identical between the sexes, suggesting that genetic architecture needs to be decoupled before SD can evolve. Furthermore, if 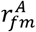 is an evolutionary important constraint, one would expect those traits with weak 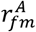 to be more likely to evolve SD, resulting in a negative relationship (Bonduriansky & Rowe, 2005; Fairbairn & Roff, 2006; Poissant *et al.*, 2010). Alternatively, selection in favor of SD may drive reductions in 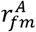, leading to the same prediction. This negative association is supported by the prevailing evidence (Poissant *et al.*, 2010), however the correlation varies widely between studies, and 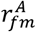 is generally a poor predictor of SD. Furthermore, 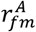 has been shown to be quickly eroded under artificial selection (Delph *et al.*, 2011).

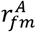 estimates provide a high-level statistical description of genotype to phenotype mapping across the sexes and are an aggregate across standing genetic variation in the population. However, we know very little about the loci that underlie this statistic. In particular, we do not know whether variation in protein coding sequence is more or less likely to cause sexually discordant phenotypic effects than expression variation. Here, we use high-throughput phenotype data from a genome-wide panel of gene knock-outs in mice to reveal unexpected differences in the gene expression architecture between the sexes (The International Mouse Phenotyping Consortium *et al.*, 2016; International Mouse Phenotyping Consortium *et al.*, 2017). We find that although most phenotypic traits are dimorphic, even many monomorphic traits harbor sex-dependent architectures, suggesting that many traits may harbor cryptic sex-specific variation. Changes in both sexes to these loci through expression may provide a way for SD to rapidly evolve, as traits are already partially decoupled and the phenotypic effect differs between males and females. These findings imply that the evolutionary constraint in SD may be more easily overcome than previously thought and explain the broad diversity of sexual dimorphism observed in nature, as well as the apparent rapid evolution of many sexually dimorphic traits.

## Results

We evaluated the sex-specific effects of alterations to gene expression, by leveraging data from large-scale high-throughput phenotyping of gene knock-out lines from the International Mouse Phenotyping Consortium (IMPC) (The International Mouse Phenotyping Consortium *et al.*, 2016). We obtained data for all continuous traits from the main IMPC pipeline for which at least 100 genotypes were available. The IMPC uses highly standardized phenotyping assays on C57BL/6 inbred mice. Both control mice and phenotype knock-out lines are tested continuously, with the eventual goal of knocking out each gene in the mouse genome. This immense scientific effort provides an unprecedented opportunity to quantify the between-sex genetic correlation across many traits and many genotypes in highly standardized conditions.

### Sexual dimorphism and 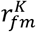 of mouse traits

If males and females share the genetic architecture of traits, knock-outs should affect the phenotype of both sexes similarly, and as architectures diverge the knock-out effects should diverge as well. We estimated the genetic correlation between males and females analogous to the conventional approach outlined above 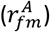. However, to delineate the knock-out lines from the traditional approach, we denote these estimates as 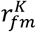, where *K* denotes the genetic variance-covariance matrix between knock-out genotypes (Figure S1). Note that 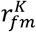 measures the correlation between the phenotypic effects of genetic knock-outs, while 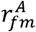 measures the correlation for genome-wide additive genetic variance.

For each of 260 traits, we obtained all available observations. On average, traits were measured in 8,069 control mice, as well as in 21,513 mice across 1,713 different knock-out genotypes. Per knock-out line, seven females and seven males were typically phenotyped.

For each trait we obtained posterior distributions for SD and the between-sex genetic correlation 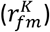 by fitting a Bayesian multilevel model. SD was expressed as the ratio of means (for Figure 1) and as the “sexual dimorphism index”: 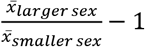 (for downstream analyses). Since mice are sexually dimorphic for body size and many traits scale with body size, we included a standardized population level effect of body weight in the model. Models without body size adjustment produced qualitatively similar results (see supplementary material). Additionally, we added group level intercepts for known sources of variance, this included the phenotyping center, the date of testing, as well as variation in testing conditions indicated by the IMPC. Using a Bayesian approach allowed us to evaluate and propagate the uncertainty in the estimate of 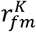 in downstream analyses. This can be important since this correlation can be biased towards 0 if it is difficult to estimate (Griffin *et al.*, 2013). Out of 260 traits tested, 202 traits passed our model evaluation procedure and were used for further inference.

**Figure 1:**
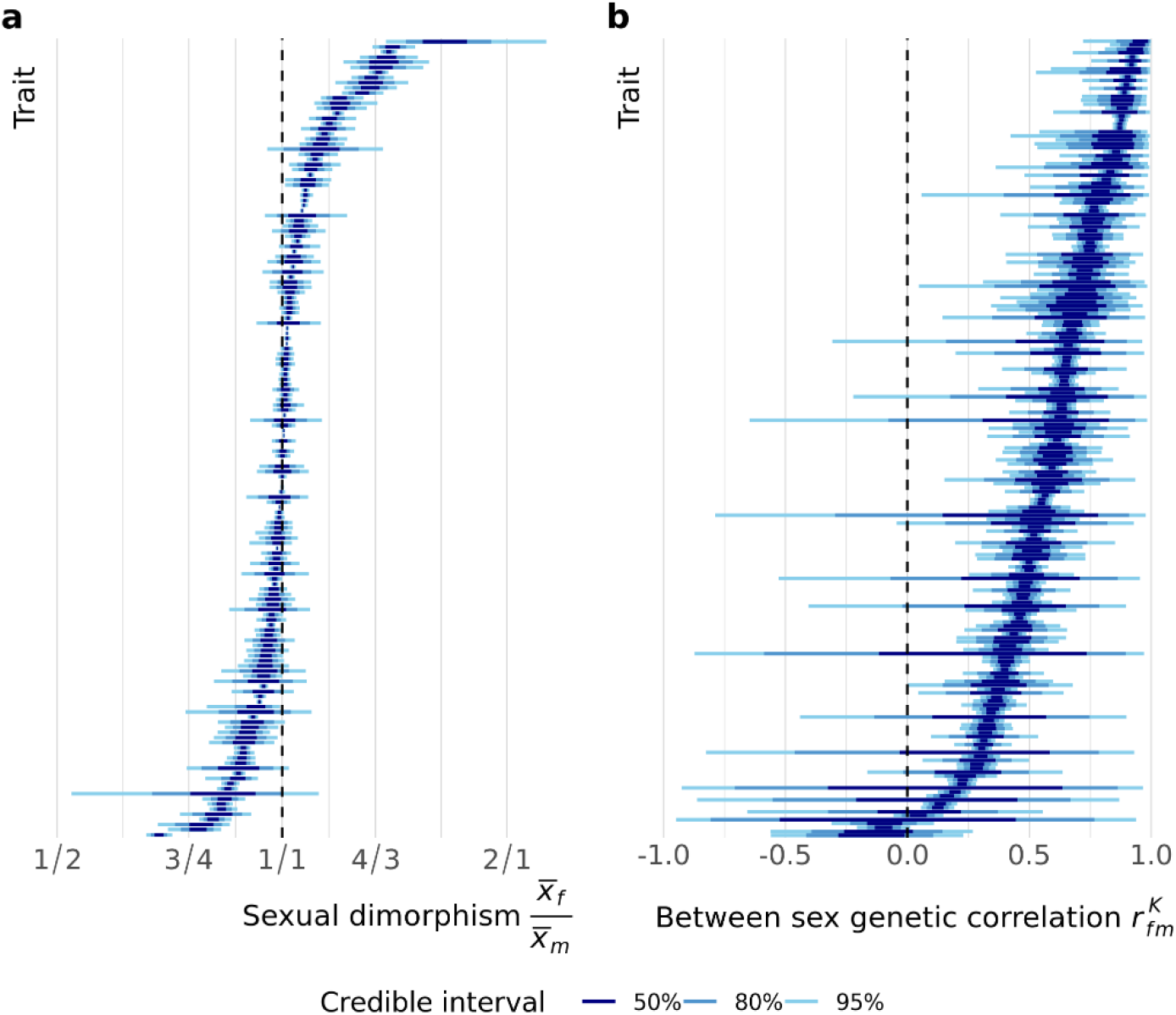
(**a**) Estimates and associated uncertainty for sexual dimorphism for each trait analyzed. Each horizontal line displays the credible intervals for one trait, where traits have been arranged by the posterior median. Shaded regions indicated the credible intervals of 50%, 80% and 95% of the posterior densities from a multilevel model. Sexual dimorphism is averaged across the wild-type genotypes, and defined as the ratio of female and male means. (**b**) As in (a), but depicting the between-sex genetic correlation 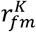. Note that the traits have been arranged independently in each panel.

Many of the measured traits showed substantial SD (Figure 1a), confirming a previous report on the IMPC data (International Mouse Phenotyping Consortium *et al.*, 2017), with an average SD index of 0.09 [0.08, 0.10] (posterior median [95% Credible Interval]). As the large sample size in this study makes it possible to distinguish small effects that have little biological relevance, we evaluated SD using equivalence testing (Wellek, 2010). We compared the 95% credible intervals (CI) of the SD index for each trait with a region of practical equivalence (ROPE) between 0 and 0.05 (Kruschke, 2018) (i.e. between 0 and 5% difference in absolute magnitude). When the entire CI falls outside the ROPE, we can be confident the sexes differ by more than 5% and the trait is considered dimorphic. We consider a trait monomorphic if we are confident there is less than a 5% difference, so when the entire CI falls within the ROPE. Under this decision rule (Kruschke, 2018), dimorphic traits roughly equal monomorphic traits. 49 out of the 156 traits (31.4%) were found to be clearly dimorphic, while 47 traits (30.1%) to be monomorphic. and 60 traits (38.5%) were not classified, as their credible interval overlapped the 5% threshold. Some of the most monomorphic traits include calcium levels in the blood and the time spent on the periphery of an open field. Strongly dimorphic traits include a variety of immune function related traits, such as spleen weight and counts of different T-cell types, as well as glucose tolerance (Table S1).

Traits showed a wide variety of estimates for 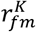, from a correlation close to 1 between the phenotypes of the sexes down to correlations indistinguishable from 0 (Figure 1b). The average correlation was clearly positive, but not as strong as we expected (0.650 [0.622, 0.689]). Surprisingly, very few traits showed a strong concordance between male and female effects, with fewer than 5% of traits having an estimate above 0.9. Some of the traits with the highest correlation are body temperature and eye morphology, while several immune phenotypes have a correlation close to 0 (Table S1).

To test the constraint that high 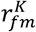 places on the evolution of dimorphism, we assessed whether 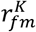 is lower for more dimorphic traits, which we would expect if dimorphism is more often associated with a reduced inter-sexual correlation. We fitted a linear model with Fisher-transformed 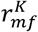 values as the dependent variable and sexual dimorphism (expressed as the SD index), propagating the uncertainty in both variables from the trait-level models. Contrary to expectation, the between-sex genetic correlation is not associated with sexual dimorphism (Figure 2, slope: −0.49 [−1.34, 0.35]). Although there is a trend in the expected direction, the relationship is non-significant, and 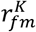 at monomorphism (i.e. the intercept) is only slightly higher than the overall average: 0.630 [0.557, 0.698].

**Figure 2:**
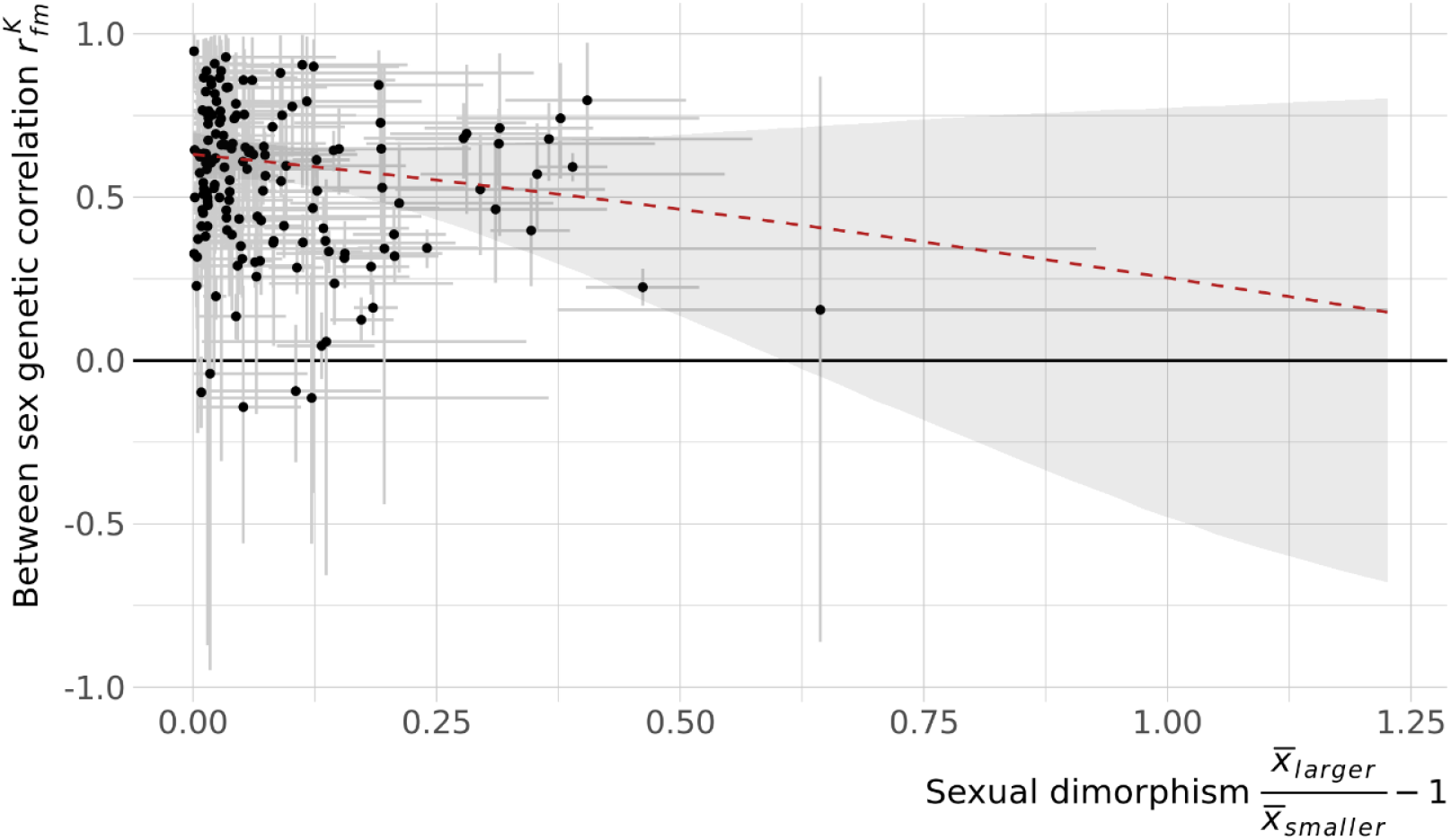
The between-sex genetic correlation does not depend on sexual dimorphism in the trait. Each point is a trait, with error bars indicating the 95% credible interval (CI) in the estimates. The red line represents the model fit of a linear model on the Fisher-transformed 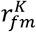, with the shaded region indicating the 95% credible interval, including propagation of trait level uncertainty. Sexual dimorphism is expressed as the SD ratio.

To investigate whether there were differences in the genetic architecture of dimorphism between trait types (Poissant *et al.*, 2010), we assigned each of the traits one of four categories: behavior, morphology, physiology or immunity (Table S1). We repeated the linear model regressing 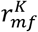 on SD, now including trait category and the SD:trait category interaction as additional parameters. There is no evidence that the relationship between 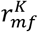 and SD is different for different trait categories (Figure S2). The average 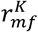 of trait categories, estimated at monomorphism, can also not clearly be distinguished (Figure S3).

Male and female genetic variances were often unbalanced, and there was a clear tendency for male genetic variance to be larger 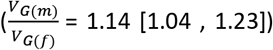. Thus, knock-out mutations have, on average, substantially larger phenotypic effects in males. It has been noted previously that mutations have larger fitness effects in male *Drosophila* (Sharp & Agrawal, 2013), and differences in genetic variance between the sexes may contribute toward the evolution of dimorphism, even under a strong between-sex genetic correlation (Wyman & Rowe, 2014). However, we found no relation between the imbalance of sex-specific variances and the level of SD (slope: 0.03 [−0.26, 0.30]).

### Development of size dimorphism and 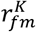

Body size is dimorphic in many species, including the mouse, yet it has been found numerous times that 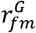 for this trait is close to 1 (Roff, 2012). Nonetheless, sexual size dimorphism can often be rapidly altered in response to the environment (Badyaev, 2002), making this an important trait to study in order to better understand the link between the evolution of SD and sex-specific architectures. As sexual size dimorphism (SSD) is established through variable development rates and times, it is especially useful to understand when in development the effect of body size loci diverges between the sexes. Unfortunately, there is very little data available for the development of 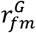, with studies usually including only 2 or 3 time points (Poissant & Coltman, 2009). In contrast, the IMPC measures body weight weekly from week 3 through 16, providing the opportunity to estimate when during development the effects of expression changes become sex-biased.

Using the same modelling approach described above, we obtained estimates for SSD and 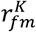 at each week (Figure 3). SSD increases strongly at the start of this period, more than doubling between weeks 3 and 7 (Figure 3a). 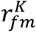 decreases during that same time (Figure 3b), and both parameters stabilize around 8 weeks. The two variables follow a roughly linear negative relationship during development (Figure 3c). A developmental link between SSD and 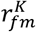 may be the result of sexually antagonistic selection mainly acting in adulthood. This would bias sex-specific loci to be expressed only later in development, driving an increasing SSD and decreasing *r_fm_*. Alternatively, strong trait integration during early development may pose significant constraints on the divergence of the sexes before 6 weeks.

**Figure 3:**
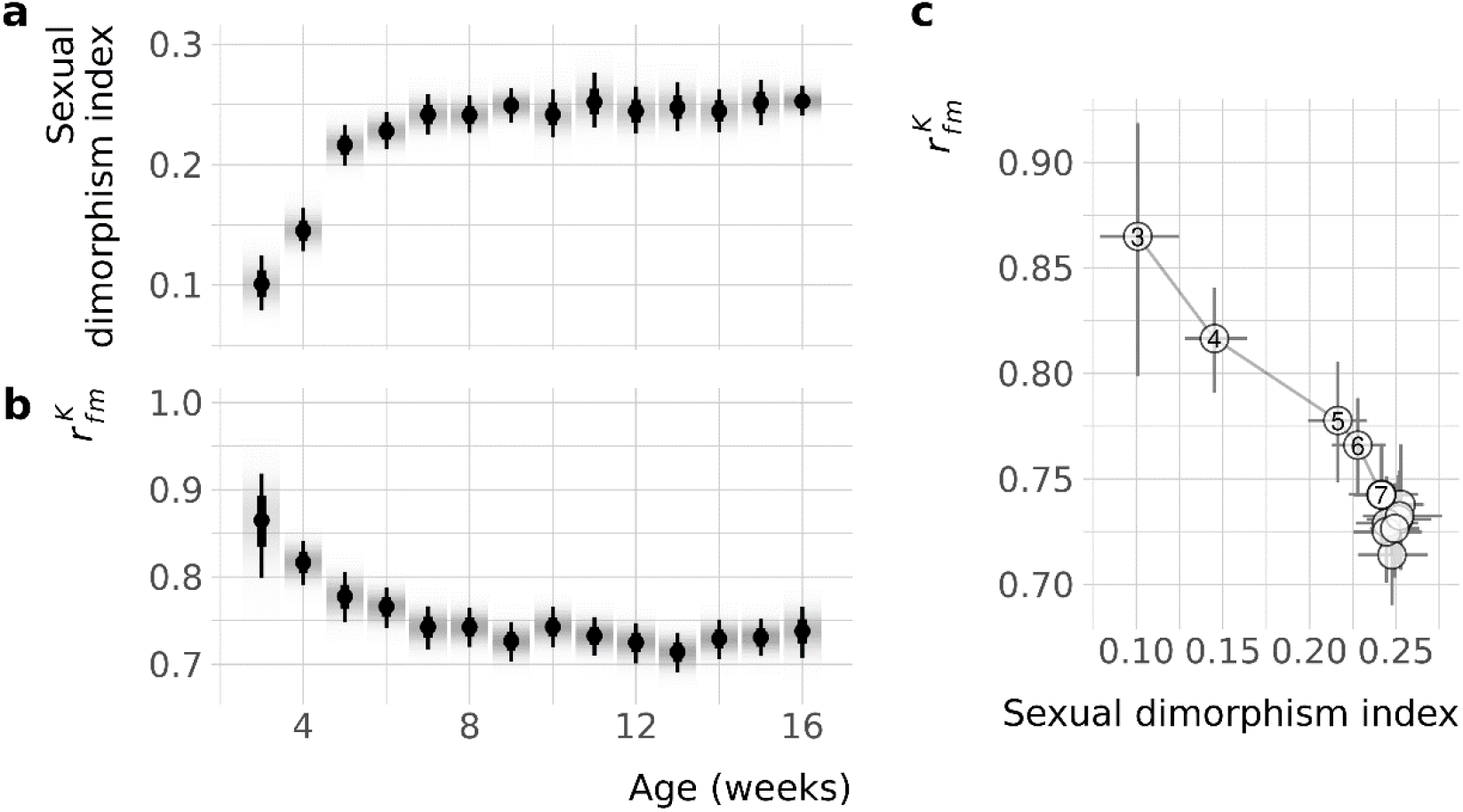
The between sex genetic correlation decreases as size dimorphism increases over development. (**a**) Estimates for sexual dimorphism in body mass for wildtype mice. Points indicate the posterior median with wide and narrow line segments denoting the 66% and 95% credible intervals respectively, and the density gradient represents the posterior density. (**b**) As in (a), but depicting the between sex genetic correlation. (**c**) Association of sexual size dimorphism and the 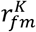 during development. Points are posterior medians with 95% credible intervals, as in (a) and (b), with lines connecting subsequent week. Weeks 3 through 7 are numbered.

### Identification of knock-out genotypes with sexually discordant effects

To gain insight into the extent to which sex-specific architectures are shared between different traits, we quantified to what extent knock-out genotypes have consistent sexually concordant or discordant effects. We separated the sexually concordant and discordant effect of each genotype on a trait by projecting the estimated effect (Best Linear Unbiased Predictor) along two independent axes (Ruzicka *et al.*, 2019), the positive and negative diagonal of a female vs male plot (as in Figure S1). Then, in order to differentiate knockouts with strong versus weak discordant effects, we looked for genotypes with a consistently low or high ranking along the discordant axis.

We identified five knock-out genotypes that consistently had smaller sexually discordant effects, compared to other genotypes (Figure 4). Those five genotypes also had much smaller concordant effects, indicating that their phenotypes are consistently average. Unsurprisingly, these were five wildtype genotypes. Additionally, 24 genotypes had larger than average discordant effects (Figure 4, Table S2). These genotypes tended to affect the sexes differently, across many traits. An analysis of Gene Ontologies for the genes that were knocked out in these genotypes, revealed no significantly overrepresented categories. In contrast to the 29 discordant genotypes, 292 genotypes (out of 2543) had consistently small or large concordant effects. This difference suggests that traits are more likely to genetically co-vary in their average value, rather than in their dimorphism.

**Figure 4:**
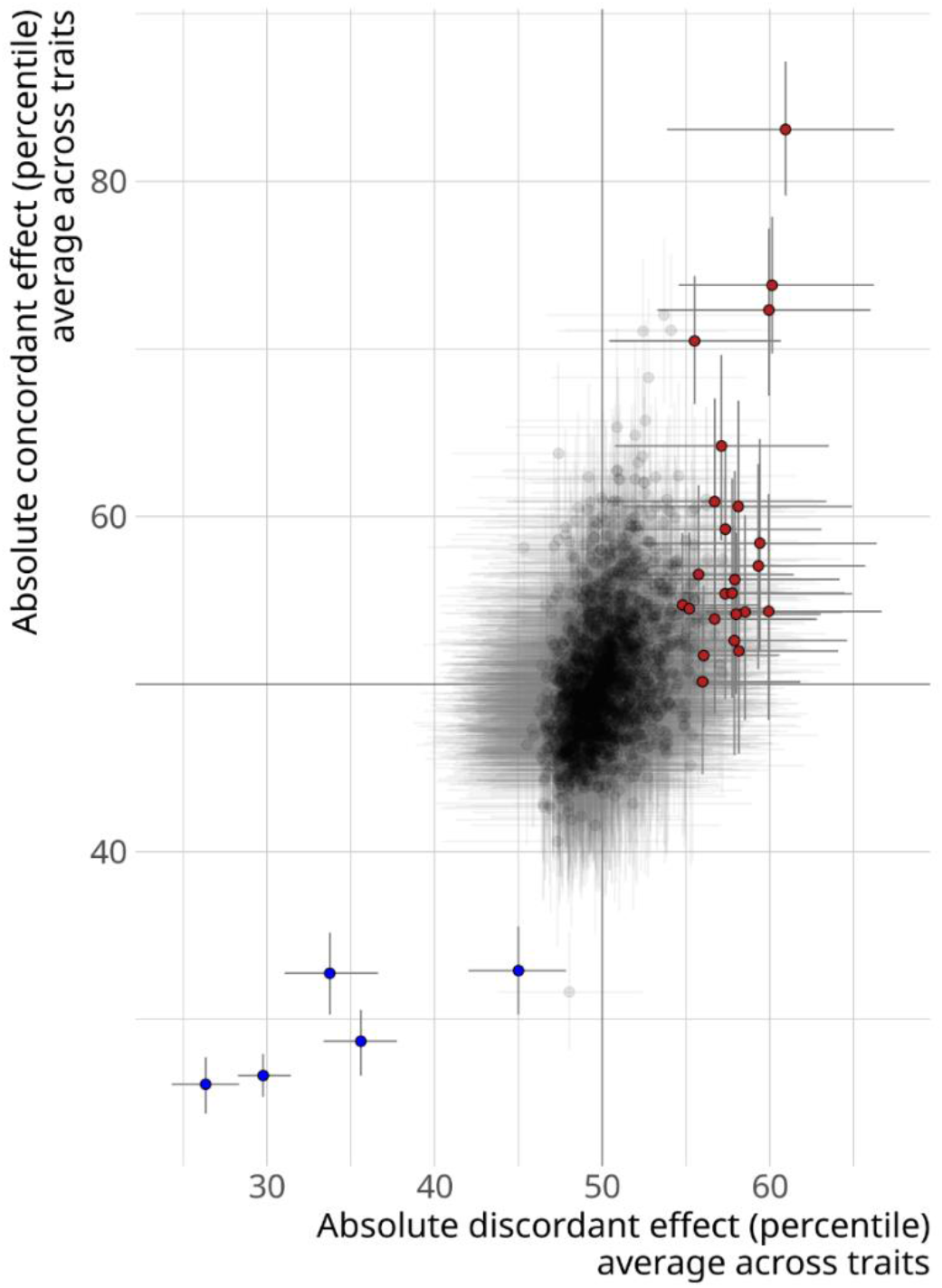
Identifying genotypes with consistent sexually discordant effects. Each point is a genotype, having been tested for at least 50 traits, with error bars denoting 95% credible intervals (CIs). The average percentile rank for the absolute sexually discordant effect of a genotype is plotted along the x-axis. The y-axis shows the average percentile tank for the absolute concordant effect. Red points indicate genotypes that tend to have more sexually discordant effects than other genotypes, while blue points are genotypes that have less discordant effects (CI does not overlap 50^th^ percentile).

### Sex-biased gene expression and fertility

Many investigations into the evolutionary significance of gene expression to SD have focused on sex-biased gene expression (Grath & Parsch, 2016). Of specific interest are expression differences in the gonads, where most sex-biased expression occurs. In these studies, it is often assumed that gonadal expression bias reflects important sex-specific fertility functions, however, it is usually not possible to verify this. Combining previously published gonadal expression data (Rinn *et al.*, 2004) with fertility data from the IMPC database, however, allowed us to test whether the expression knock-out of sex-biased genes causes sex-specific infertility.

As predicted, fertility status was significantly associated with expression bias category (i.e. male-biased, female-biased or unbiased; χ^2^_6_ = 76.6, p < 0.001, Figure S4). Gene knockouts of female-biased or unbiased genes led to male-limited infertility in 1.5% of cases, but this increased to 11% of cases when knocking out male-biased genes. Female-limited fertility on the other hand was less common in general and showed no increase with knock-outs of female-biased genes (Figure S4), possibly because female gametogenesis is largely encoded during fetal development and then arrested.

## Discussion

Using the extensive phenotyping effort of gene knock-out mouse lines by the IMPC, we have tested for the extent of overlap in trait genetic architecture between males and females. Even in the mouse, which is relatively monomorphic when compared to many other vertebrates, it is surprisingly common for traits to show clear differences between the sexes after controlling for body size. This therefore suggests that sexual dimorphism is not the exception but the norm across many crucial somatic traits.

Furthermore, traits are affected differently by knock-out mutations depending on the sex of the individual. This clearly illustrates that studies of gene function must account for sex, as knock-out effects may only be easily detectable in one of the sexes (International Mouse Phenotyping Consortium *et al.*, 2017). Alterations in gene expression are often thought to be a common mechanism to resolve intra-locus sexual conflict by making gene expression sex-biased or sex-specific (Grath & Parsch, 2016). This assumes a shared genetic architecture, which is differentially regulated between the sexes. Our work suggests that the underlying architecture may differ between the sexes in many cases, and the low estimates of 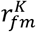 that we recover highlight a different potential role of gene expression in the evolution of SD.

Mutations of large regulatory effect can often be expected to alter SD, providing one way to resolve intra-locus sexual conflict. However, these regulatory changes need not result in sex-biased gene expression, as our work suggests that regulatory changes in both sexes, in this case elimination of expression in both sexes through knockouts, often predominantly only affect the phenotype of one. In other words, sexually concordant regulatory changes can result in sexually discordant phenotypic effects, and our results suggest that this commonly occurs. This provides an alternative route to dimorphism through sex-specific genetic architecture, rather than differential regulation of shared architecture. This could, for example, be the result of interactions with sex-biased genes in the same regulatory network, or of a sex-bias in the size of the cell populations expressing the gene. It appears likely that the modulation of gene expression, either through sex-bias in the downstream phenotypic effects or in the expression itself, is a major contributor to the evolution of SD.

Although mutations of large effect, especially gene deletions, can have deleterious effects on other traits through pleiotropy, many genes are non-essential (Amsterdam *et al.*, 2004; Liao & Zhang, 2007; Georgi *et al.*, 2013). This suggests significant regulatory potential in the evolution of SD. Additionally, the knockout mutations assessed here likely represent an extreme form of regulatory variation, which we would expect to have similar, if less drastic, sex-specific effects, and more often contribute to SD.

As others have previously indicated (Cowley & Atchley, 1988; Reeve & Fairbairn, 2001; Bonduriansky & Rowe, 2005), 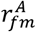 may not be as strong an indicator of constraint as was originally suggested (Lande, 1980). While 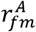 is very useful in describing the potential for the standing genetic variation to alter SD in a single or a few generations, it cannot detect decoupling in trait architectures which are currently lacking variation. Our results indicate that even high 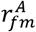 traits may be susceptible to changes in SD, as most traits have cryptic parts of the genetic architecture in which new mutations will likely have sex discordant effects. Importantly, changes in the architecture itself, such as changes in gene pathways or the recruitment of new transcription factors, are not necessary to have occurred, contrasting with a common interpretation of a strong 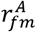.

A potential limitation of this study is that the mice are inbred, resulting in genome-wide homozygosity. This means that the phenotypic variation is expected to be relatively small, making the effects of knockouts appear stronger. Additionally, the effects of dominance and epistasis are artificially limited. As it has been suggested that sex-specific dominance is pervasive (Grieshop & Arnqvist, 2018), and epistatic interactions could be affected by sex as well, our estimates of 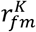 could potentially be biased upwards. It is also important to note that sex-linked genetic architecture can allow for the evolution of dimorphism. However, given the relatively small size and limited gene content of the mouse Y chromosome (Soh *et al.*, 2014), the role of the Y in sex-specific genetic architecture for a broad array of somatic traits is unclear.

The vast majority of genotypes were neither strongly nor weakly discordant across traits, suggesting there are very few or no “sex-specific genes” or “SD genes” but rather many different genes have sex-specific effects on different traits. The few genotypes that did show some consistently discordant effects had no functional categories in common, also suggesting that SD is regulated differently in different traits. As we identified more genotypes that had consistently large concordant effects, the genetic covariance between trait means is likely stronger than between SD of different traits. Large-scale analyses in a multivariate framework are needed to fully clarify the covariance of expression variance across traits and sex, in order to come to a complete understanding of the evolutionary constraints on SD.

In conclusion, using a dataset of unprecedented size, we have demonstrated that traits harbor a surprising amount of sex-specific genetic architecture, as sexes respond variably to knock-out mutations. These results may help explain why SD is common, evolvable and variable, even under supposed strong genetic constraints. While these differences clearly indicate that the genotype-to-phenotype mapping is sex-dependent for most traits, it remains unclear what underlying mechanisms are the cause for this. We hope future work will help elucidate proximate causes and evolutionary consequences of this work.

## Methods

We obtained data from the online IMPC genotype-phenotype database. We selected phenotypes for analysis by requesting all uni-dimensional continuous traits, excluding legacy pipelines. We also excluded traits that were not measured in both sexes, fitness-related traits (such as reproductive screening), body size (we analyzed body size separately), traits with fewer than 100 genotypes, and traits that were clearly not actually continuous (such as a count of the number of ribs). After triage, we had 260 traits for which we downloaded all available phenotype data, including both knock-out phenotypes and control data. On average we obtained data for 8,069 control mice and 21,513 mice from 1,713 knock-out lines, per trait.

### Sexual dimorphism and 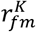 of mouse traits

As we were interested in estimating a single value for 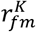 per trait, we collapsed different sources of genetic variance into genotypes. As some gene knock-outs were performed in different genetic backgrounds, some genes had multiple allelic knock-outs, and some genes were tested in different zygosities, we defined each unique gene:allele:background:zygosity combination as a separate genotype. Note that the genetic backgrounds are all C57BL/6 mice, but a different sub-strain.

To each of the trait datasets, we fitted a Bayesian linear mixed model with the goal of estimating both the between-sex genetic correlation 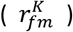 and sexual dimorphism (SD). We opted for the analysis of single traits as opposed to multivariate models, since phenotypes have been measured across differing sets of individuals and knock-outs. Additionally, the univariate models were computationally expensive, with each model taking several days to a week to fit, and multivariate models would be logistically unfeasible. Each model had one of the phenotypes as the dependent variable, which was standardized (centered and scaled to unit variance) and transformed (see below). We included sex as a population level effect (also called fixed effect), allowing an average level of dimorphism across genotypes, although we did not directly use this parameter as our measurement of SD (see below). We also included body mass as a population level parameter, since mice are size dimorphic. Body mass was standardized (centered and scaled to unit variance) prior to analysis. All analyses were repeated without body mass, and the qualitatively similar results can be found in the supplementary material, although we only recommend interpretation of the results accounting for body size.

To estimate 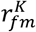 we added group level parameters (also called random effects) of genotype for each sex, and their correlation. Finally, we added group level intercepts for known sources of variation when they were present, which were 1) the phenotyping center in which testing was performed, a parameter encoding several methodological differences (“meta group”), and 2) the date of testing. This leads to the final model definition (in *lme4/brms* syntax): *phenotype ~ weight + sex + (0 + sex | genotype) + (1 | center) + (1 | meta_group) + (1 | date)*. In mathematical notation, following Gelman & Hill (2006):

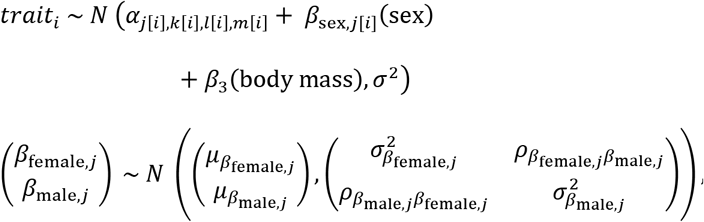

for genotype j = 1,…, J

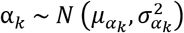, for center k = 1,…, K

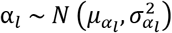, for meta group l = 1,…, L

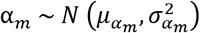, for date m = 1,…, M

Parameter values were estimated using the brms (Bürkner, 2017, 2018) interface to the probabilistic programming language Stan (Carpenter et al., 2017). We used weakly informative prior distributions, with priors of N(0, 1) for the intercept and N(0, 2) for the effect of body mass. For the group level standard deviations and residual standard deviation we used the positive range of unit student-t distributions with 5 degrees of freedom. Finally, we used an LKJ prior with η = 1 for 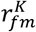, which is uniform over the range −1 to 1. Posterior distributions were obtained using Stan’s no-U-turn HMC sampler, with 2 chains of 8000 iterations, with the first 4000 used as warm-up and discarded. We additionally set the max tree-depth to 20 and the adapt delta parameter to 0.9. To evaluate the ability of our models to accurately estimate the between-sex genetic correlation, even though the sample size for each genotype was limited, we performed a simulation study (figure S7), confirming that our approach recovers the true value for 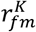.

In order to satisfy the assumption of approximately normal residuals, we preceded each analysis by estimation of a Box-Cox transformation, following the established methods by the IMPC (Kurbatova *et al.*, 2019), using the simplified model definition: *phenotype ~ weight + sex + (0 + sex | genotype)*. We estimated the transform using the *bcnPower* method in the *car* package (Fox *et al.*, 2019), with model fitting performed by *lme4* (Bates *et al.*, 2015).

After fitting all 260 trait models, we performed model criticism. For each model we obtained the maximum 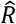 parameter, the number of divergences and the minimum effective sample size. We removed all models that had a maximum 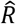 of more than 1.05, more than 2.5% divergent draws, or a minimum effective sample size of less than 400. Finally, we performed visual posterior predictive checks (Gabry *et al.*, 2019), and removed models that did not reproduce the observed data distribution. Considering the computational effort required for each of these models, as well as that the number of successful models was more than large enough for the analyses we wished to perform, we did not attempt to remedy the failing models. We performed visual checks to confirm that the excluded traits did not have a bias in SD or 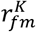. After model criticism, 202 out of 260 models remained.

For each of these models we derived posterior distributions of 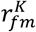. Note that *brms* estimates standard deviations and correlations directly, so no parameter transformation was necessary. We then derived posterior distributions of SD by predicting average male and female phenotypes for wildtype (i.e. control group) mice. When there were multiple genetic background variations in which a trait was tested, we used the marginal means across backgrounds. To make SD estimates comparable across traits, we used a mean standardized effect size for SD, the SD index: 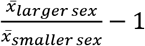, i.e. the ratio between larger and smaller, divided by the residual standard deviation. Note that the SD index requires that comparisons to zero are biologically meaningful (i.e. traits are measured on a ratio scale), which was not true for all the traits in our data set, such as body temperature, indices and fractional measures. We therefore performed back transformations of the marginal means to the original scale, and we only calculated SD for 156 out of 202 traits.

After obtaining the posteriors for each trait, we used a linear model to test for a relationship between 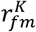 and SD. In order to account for uncertainty in those estimates we performed random draws from the posterior distributions of those estimates to create 500 datasets. For each of those samples we ran one MCMC chain of a 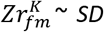 *index* model using the *brm_multiple* function, and performed inference on the combined set of 500 chains. Note that we performed a Z-transformation on 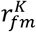, also called the Fisher transformation, to stabilize the variance. Additionally, we performed the same procedure for the ratio of the genetic variances: 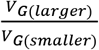, which was log transformed before analysis.

### Development of size dimorphism and 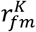

To quantify sexual size dimorphism during development, and associated changes in 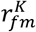, we split the body mass data into different ages. Mice were weighed once a week, with most mice being measured between 4 and 16 weeks of age. For each week, we ran the same analysis as for the separate traits outlined above.

### Identification of knock-out genotypes with sexually discordant effects

The concordant and discordant nature of knock-out genotypes was determined by evaluating whether the genotypes were consistently ranked low or high along the concordant and discordant axes across traits. For each trait, we used the multilevel model that was used to estimate SD and 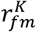, described above, to obtain estimates of the male and female trait values for the measured genotypes. We extracted the posteriors for the male and female parameter for the genotype group term (BLUP). Note that these estimates are adjusted for body weight and environmental effects, have already undergone parameter shrinkage, and are centered around zero. We then translated the male and female phenotypes into concordant and discordant effects, by rotating the axes so that the concordant axis is the positive diagonal (female = male) and the discordant axis is the negative diagonal (female = -male). The absolute value along the two diagonal axes was taken, so that the effect of a genotype is larger when it is further from the population average. Since the size of the discordant effects of a genotype is strongly affected by the trait architecture (i.e. 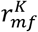), we assigned genotypes percentile ranks to aid comparison across traits.

For all genotypes that were tested for at least 100 phenotypes, we then calculated the average concordant and discordant rank across traits. Credible intervals (CIs) for this average were calculated by computing that average for 100 random draws of the posteriors. We then categorized genotypes as less or more discordant than average by checking whether the CI overlapped a median rank (50^th^ percentile in Figure 4).

For the genotypes that were more discordant than average, we extracted which gene had been knocked out and analyzed the associated gene ontology (GO) terms. Using *goseq* (Young *et al.*, 2010) we tested for overrepresented GO terms, using the hypergeometric method for obtaining p-values. Finally, we adjusted the p-values to control the false discovery rate(Benjamini & Hochberg, 1995).

### Sex-biased gene expression and fertility

We obtained published gene expression profiles of male and female gonadal tissue from the ArrayExpress database under accession number E-GEOD-1148 (Rinn *et al.*, 2004). Using *limma* (Ritchie *et al.*, 2015), we calculated the difference in expression between the sexes (log_2_ fold change), and empirical Bayes moderated t-statistics with adjusted p-values. We then classified genes as sex-biased if the fold-change was at least 2, and the adjusted p-values was significant (α = 0.05). Genes that did not satisfy both those criteria were categorized as unbiased.

We then obtained female and male specific fertility data from the IMPC (phenotypes IMPC_FER_019_001 and IMPC_FER_001_001), which are binary traits (fertile vs. infertile) where each sex has been allowed to breed with a wildtype mate. Combining these we defined four fertility categories: fertile, female-limited infertile, male-limited infertile and infertile. To test for an association between gene expression category and fertility outcome after knock-out, we performed a 3×4 chi-squared test for independence.

#### Software

All analyses were performed in R v3.6.1 (R Core Team, 2019). Specific R packages used in the analyses are listed above, and the *tidyverse* (Wickham *et al.*, 2019) was used for general data handling and visualization.

## Acknowledgements

We gratefully acknowledge support from a Canada 150 Research Chair and the European Research Council (grant agreement 680951). We thank Locke Rowe, Michael Whitlock and members of the Mank Lab for helpful comments and suggestions. Finally, we are indebted to all contributors to the IMPC project and their commitment to open science.

## Author Contributions

Both authors conceived of the study, WvdB performed the data analysis, and both authors wrote the manuscript.

## Data Availability Statement

No new data was collected for this study. All raw phenotype data is available from the International Mouse Phenotyping Consortium (https://www.mousephenotype.org/). The gene expression profiles of male and female gonadal tissue is available from the ArrayExpress database under accession number E-GEOD-1148. All estimates used in down-stream analyses are available in the Supplementary Information.

## References

Amsterdam, A., Nissen, R.M., Sun, Z., Swindell, E.C., Farrington, S. & Hopkins, N. 2004. Identification of 315 Genes Essential for Early Zebrafish Development. Proc. Natl. Acad. Sci. U. S. A. 101: 12792–12797.

Bachtrog, D., Mank, J.E., Peichel, C.L., Kirkpatrick, M., Otto, S.P., Ashman, T.-L., et al. 2014. Sex Determination: Why So Many Ways of Doing It? PLoS Biol. 12: e1001899.

Badyaev, A.V. 2002. Growing apart: an ontogenetic perspective on the evolution of sexual size dimorphism. Trends Ecol. Evol. 17: 369–378.

Bates, D., Mächler, M., Bolker, B. & Walker, S. 2015. Fitting Linear Mixed-Effects Models Using lme4. J. Stat. Softw. 67: 1–48.

Benjamini, Y. & Hochberg, Y. 1995. Controlling the False Discovery Rate: A Practical and Powerful Approach to Multiple Testing. J. R. Stat. Soc. Ser. B Methodol. 57: 289–300.

Bonduriansky, R. & Rowe, L. 2005. Intralocus Sexual Conflict and the Genetic Architecture of Sexually Dimorphic Traits in Prochyliza Xanthostoma (diptera: Piophilidae). Evolution 59: 1965–1975.

Bürkner, P.-C. 2018. Advanced Bayesian Multilevel Modeling with the R Package brms. R J. 10: 395–411.

Bürkner, P.-C. 2017. brms: An R Package for Bayesian Multilevel Models Using Stan. J. Stat. Softw. 80: 1–28.

Carpenter, B., Gelman, A., Hoffman, M.D., Lee, D., Goodrich, B., Betancourt, M., et al. 2017. Stan: A Probabilistic Programming Language. J. Stat. Softw. 76.

Chenoweth, S.F., Rundle, H.D. & Blows, M.W. 2008. Genetic Constraints and the Evolution of Display Trait Sexual Dimorphism by Natural and Sexual Selection. Am. Nat. 171: 22–34.

Cheverud, J.M., Vaughn, T.T., Pletscher, L.S., Peripato, A.C., Adams, E.S., Erikson, C.F., et al. 2001. Genetic architecture of adiposity in the cross of LG/J and SM/J inbred mice. Mamm. Genome 12: 3–12.

Cowley, D.E. & Atchley, W.R. 1988. Quantitative Genetics of Drosophila Melanogaster. II. Heritabilities and Genetic Correlations between Sexes for Head and Thorax Traits. Genetics 119: 421–433.

Cowley, D.E., Atchley, W.R. & Rutledge, J.J. 1986. Quantitative Genetics of Drosophila Melanogaster. I. Sexual Dimorphism in Genetic Parameters for Wing Traits. Genetics 114: 549–566.

Delph, L.F., Steven, J.C., Anderson, I.A., Herlihy, C.R. & Iii, E.D.B. 2011. Elimination of a Genetic Correlation Between the Sexes Via Artificial Correlational Selection. Evolution 65: 2872–2880.

Ellegren, H. & Parsch, J. 2007. The evolution of sex-biased genes and sex-biased gene expression. Nat. Rev. Genet. 8: 689–698.

Fairbairn, D.J. & Roff, D.A. 2006. The quantitative genetics of sexual dimorphism: assessing the importance of sex-linkage. Heredity 97: 319–328.

Fox, J., Weisberg, S., Price, B., Adler, D., Bates, D., Baud-Bovy, G., et al. 2019. car: Companion to Applied Regression.

Gabry, J., Simpson, D., Vehtari, A., Betancourt, M. & Gelman, A. 2019. Visualization in Bayesian workflow. J. R. Stat. Soc. Ser. A Stat. Soc. 182: 389–402.

Galouzis, C.C. & Prud’homme, B. 2021. Transvection regulates the sex-biased expression of a fly X-linked gene. Science 371: 396–400. American Association for the Advancement of Science.

Gelman, A. & Hill, J. 2006. Data Analysis Using Regression and Multilevel/Hierarchical Models. Cambridge University Press.

Georgi, B., Voight, B.F. & Bucan, M. 2013. From mouse to human: evolutionary genomics analysis of human orthologs of essential genes. PLoS Genet. 9.

Grath, S. & Parsch, J. 2016. Sex-Biased Gene Expression. Annu. Rev. Genet. 50: 29–44.

Grieshop, K. & Arnqvist, G. 2018. Sex-specific dominance reversal of genetic variation for fitness. PLOS Biol. 16: e2006810.

Griffin, R.M., Dean, R., Grace, J.L., Ryden, P. & Friberg, U. 2013. The Shared Genome Is a Pervasive Constraint on the Evolution of Sex-Biased Gene Expression. Mol. Biol. Evol. 30: 2168–2176.

Hanrahan, J.P. & Eisen, E.J. 1973. Sexual dimorphism and direct and maternal genetic effects on body weight in mice. Theor. Appl. Genet. 43: 39–45.

Hansen, T.F. 2006. The Evolution of Genetic Architecture. Annu. Rev. Ecol. Evol. Syst. 37: 123–157.

Hermansen, J.S., Starrfelt, J., Voje, K.L. & Stenseth, N.C. 2018. Macroevolutionary consequences of sexual conflict. Biol. Lett. 14: 20180186.

International Mouse Phenotyping Consortium, Karp, N.A., Mason, J., Beaudet, A.L., Benjamini, Y., Bower, L., et al. 2017. Prevalence of sexual dimorphism in mammalian phenotypic traits. Nat. Commun. 8.

Khila, A., Abouheif, E. & Rowe, L. 2012. Function, Developmental Genetics, and Fitness Consequences of a Sexually Antagonistic Trait. Science 336: 585–589. American Association for the Advancement of Science.

Kruschke, J.K. 2018. Rejecting or Accepting Parameter Values in Bayesian Estimation. Adv. Methods Pract. Psychol. Sci. 1: 270–280.

Kurbatova, N., Karp, N., Mason, J. & Haselimashhadi, H. 2019. PhenStat: Statistical analysis of phenotypic data. Bioconductor version: Release (3.10).

Lande, R. 1980. Sexual dimorphism, sexual selection and adaptation in polygenic characters. Evolution 34: 292–305.

Liao, B.-Y. & Zhang, J. 2007. Mouse duplicate genes are as essential as singletons. Trends Genet. 23: 378–381.

Mank, J.E. 2017. The transcriptional architecture of phenotypic dimorphism. Nat. Ecol. Evol. 1: 1–7. Nature Publishing Group.

Owens, I.P.F. & Hartley, I.R. 1998. Sexual dimorphism in birds: why are there so many different forms of dimorphism? Proc. R. Soc. Lond. B Biol. Sci. 265: 397–407.

Poissant, J. & Coltman, D.W. 2009. The ontogeny of cross-sex genetic correlations: an analysis of patterns. J. Evol. Biol. 22: 2558–2562.

Poissant, J., Wilson, A.J. & Coltman, D.W. 2010. Sex-specific genetic variance and the evoltuion of sexual dimorphism: a systematic review of cross-sex genetic correlations. Evolution 64: 97–107.

R Core Team. 2019. R: A Language and Environment for Statistical Computing. R Foundation for Statistical Computing, Vienna, Austria.

Reeve, J.P. & Fairbairn, D.J. 2001. Predicting the evolution of sexual size dimorphism. J. Evol. Biol. 14: 244–254.

Rice, W.R. & Chippindale, A.K. 2001. Intersexual ontogenetic conflict. J. Evol. Biol. 14: 685–693.

Rinn, J.L., Rozowsky, J.S., Laurenzi, I.J., Petersen, P.H., Zou, K., Zhong, W., et al. 2004. Major Molecular Differences between Mammalian Sexes Are Involved in Drug Metabolism and Renal Function. Dev. Cell 6: 791–800.

Ritchie, M.E., Phipson, B., Wu, D., Hu, Y., Law, C.W., Shi, W., et al. 2015. limma powers differential expression analyses for RNA-sequencing and microarray studies. Nucleic Acids Res. 43: e47–e47.

Roff, D.A. 2012. Evolutionary Quantitative Genetics. Springer Science & Business Media.

Ruzicka, F., Hill, M.S., Pennell, T.M., Flis, I., Ingleby, F.C., Mott, R., et al. 2019. Genome-wide sexually antagonistic variants reveal long-standing constraints on sexual dimorphism in fruit flies. PLOS Biol. 17: e3000244.

Sharp, N.P. & Agrawal, A.F. 2013. Male-biased fitness effects of spontaneous mutation in drosophila melanogaster. Evolution 67: 1189–1195.

Soh, Y.Q.S., Alföldi, J., Pyntikova, T., Brown, L.G., Graves, T., Minx, P.J., et al. 2014. Sequencing the Mouse Y Chromosome Reveals Convergent Gene Acquisition and Amplification on Both Sex Chromosomes. Cell 159: 800–813.

Stewart, A.D. & Rice, W.R. 2018. Arrest of sex-specific adaptation during the evolution of sexual dimorphism in Drosophila. Nat. Ecol. Evol. 2: 1507–1513.

The International Mouse Phenotyping Consortium, Dickinson, M.E., Flenniken, A.M., Ji, X., Teboul, L., Wong, M.D., et al. 2016. High-throughput discovery of novel developmental phenotypes. Nature 537: 508–514.

Wellek, S. 2010. Testing statistical hypotheses of equivalence and noninferiority. Chapman and Hall/CRC.

Wickham, H., Averick, M., Bryan, J., Chang, W., McGowan, L., François, R., et al. 2019. Welcome to the Tidyverse. J. Open Source Softw. 4: 1686.

Wyman, M.J. & Rowe, L. 2014. Male Bias in Distributions of Additive Genetic, Residual, and Phenotypic Variances of Shared Traits. Am. Nat. 184: 326–337.

Young, M.D., Wakefield, M.J., Smyth, G.K. & Oshlack, A. 2010. Gene ontology analysis for RNA-seq: accounting for selection bias. Genome Biol. 11: R14.

