## Supplementary information for "Widespread cryptic variation in genetic architecture between the sexes"

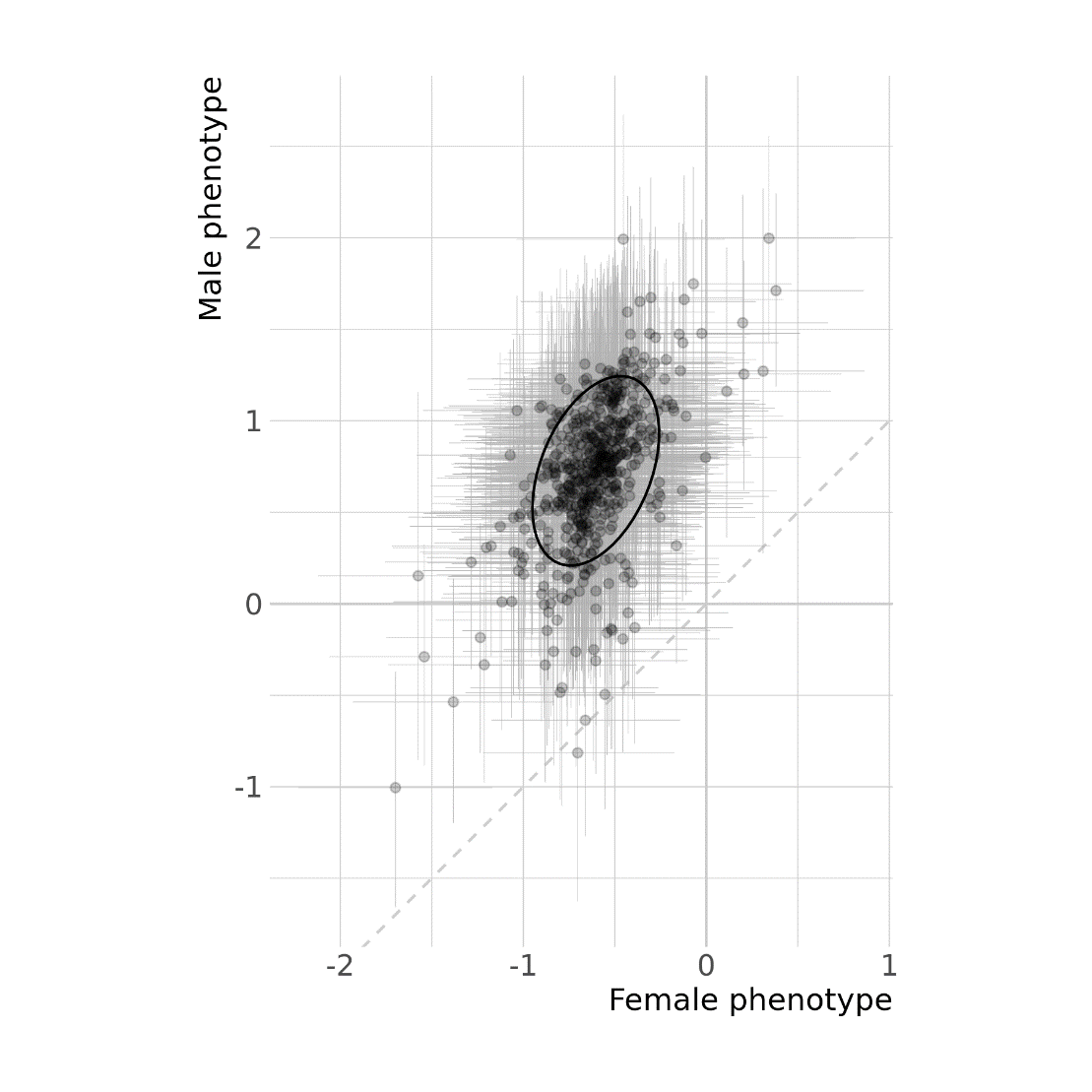

**Figure S1**: Analysis of sex-specific genetic variance between IMPC knock-out lines, using spleen weight as an example. Each point represents an estimate (best linear unbiased predictor, BLUP) of the male and female phenotypes of one knock-out genotype, estimated from a Bayesian mixed model accounting for body weight and other factors. Spleen weight was Box-Cox transformed and standardized before analysis. The ellipse drawn describes the observed genetic variance-covariance matrix, and the diagonal dotted line signifies monomorphism. $r_{fm}^{K}$ for this trait is 0.40 (posterior median, 95% credible interval: [0.23, 0.56]).

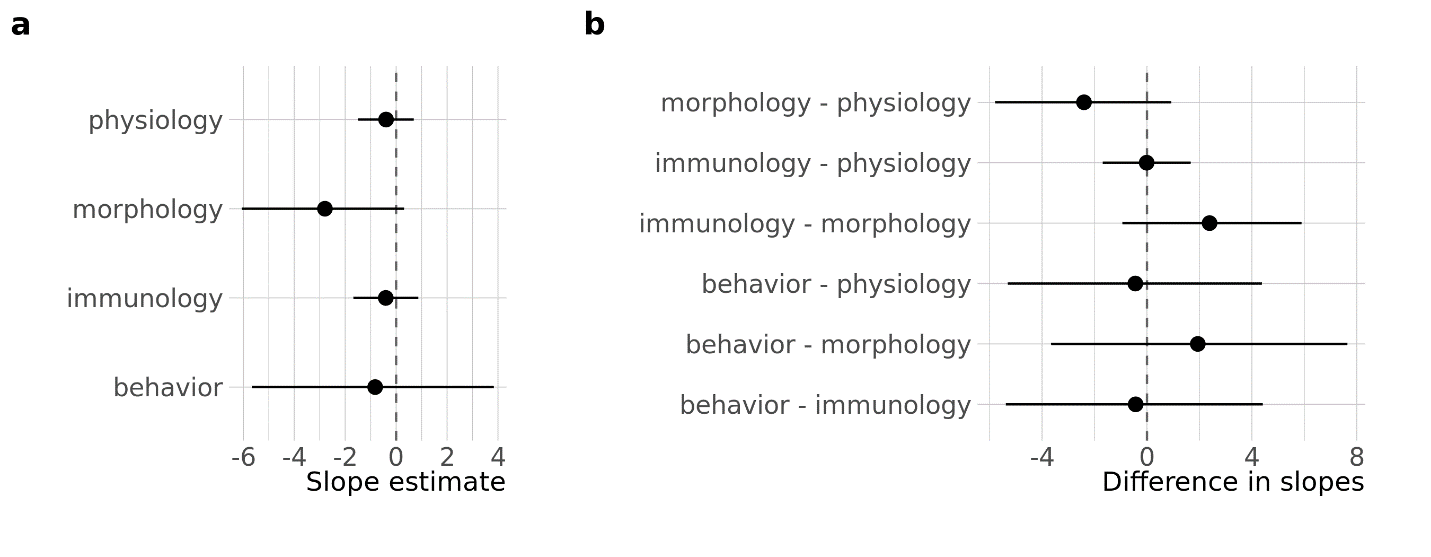

**Figure S2:** Relationship between sexual dimorphism and the between sex genetic correlation $r_{mf}^{K}$. Panel **a** shows the marginal estimates for the slope (${Zr}_{mf}^{K}\sim SD$) for each trait category. Panel **b** shows the contrasts between the slopes.

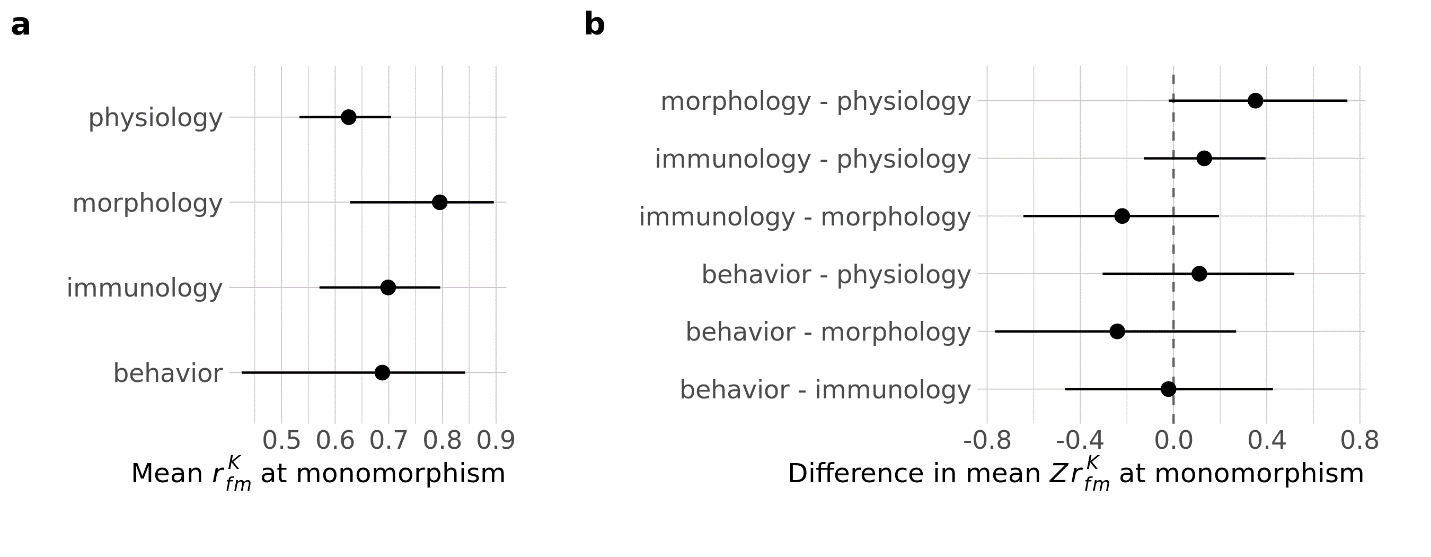

**Figure S3:** Comparison of $r_{mf}^{K}$ at monomorphism (the model intercept) between trait categories. Panel **a** shows the marginal means. Panel **b** shows the contrasts between the means.

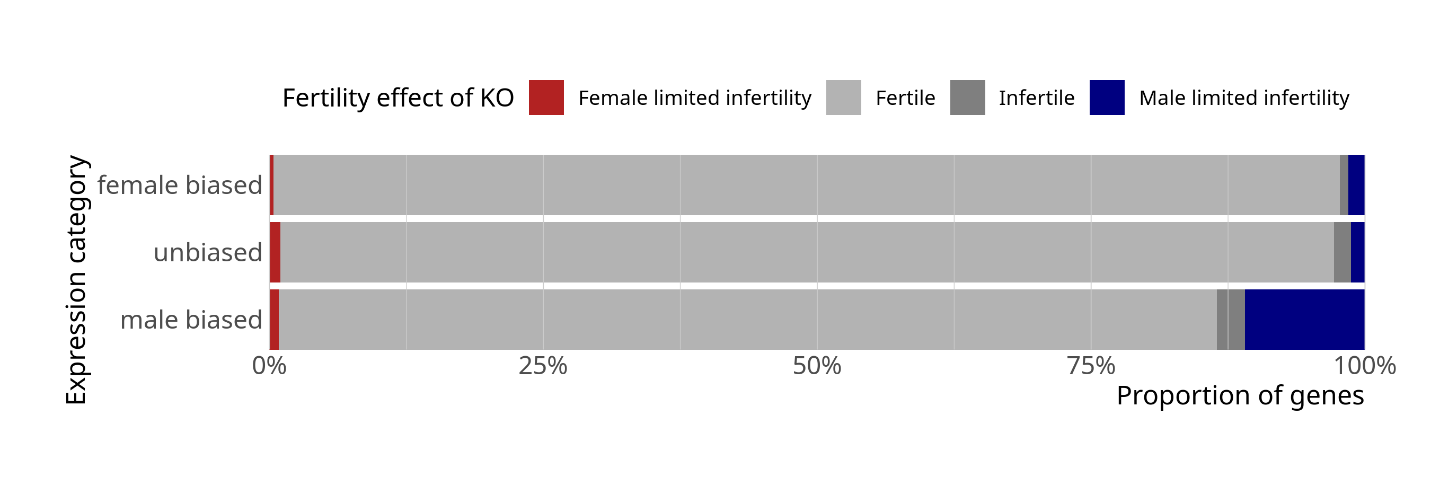

**Figure S4:** Fertility of gene knock-out lines, comparing between genes that have sex-biased expression in the gonads, and genes that have unbiased expression.

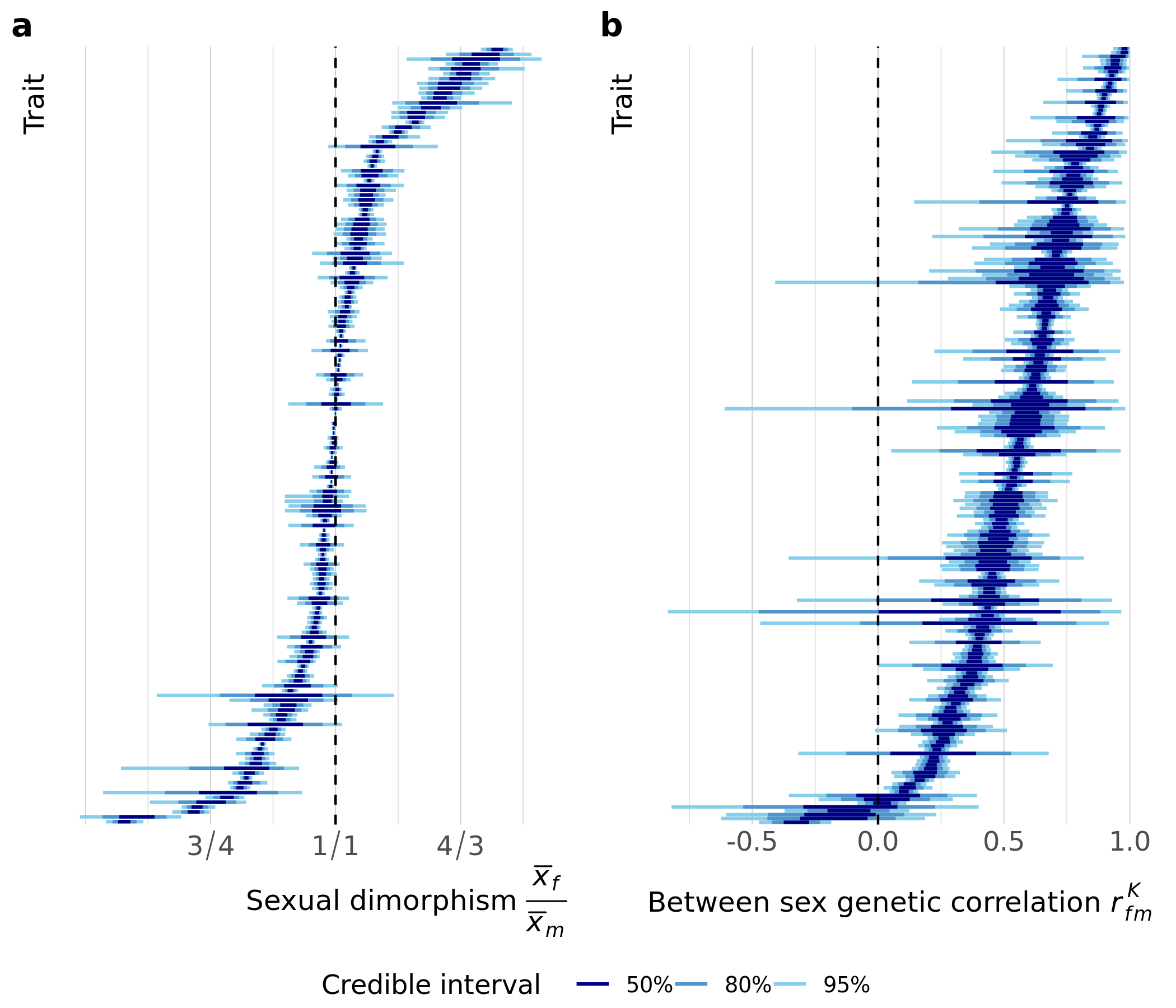

**Figure S5:** As Figure 1, but without accounting for body mass. (**a**) Estimates and associated uncertainty for sexual dimorphism for each trait analyzed. Each line displays the credible intervals for one trait, where traits have been arranged by the posterior median. Shaded regions indicated the credible intervals of 50%, 80% and 95% of the posterior densities from a multi-level model. Sexual dimorphism is averaged across the wild-type genotypes, and defined as the ratio of female and male means. (**b**) As in (a), but depicting the between sex genetic correlation $r_{fm}^{K}$. Note that the traits have been arranged independently in each panel.

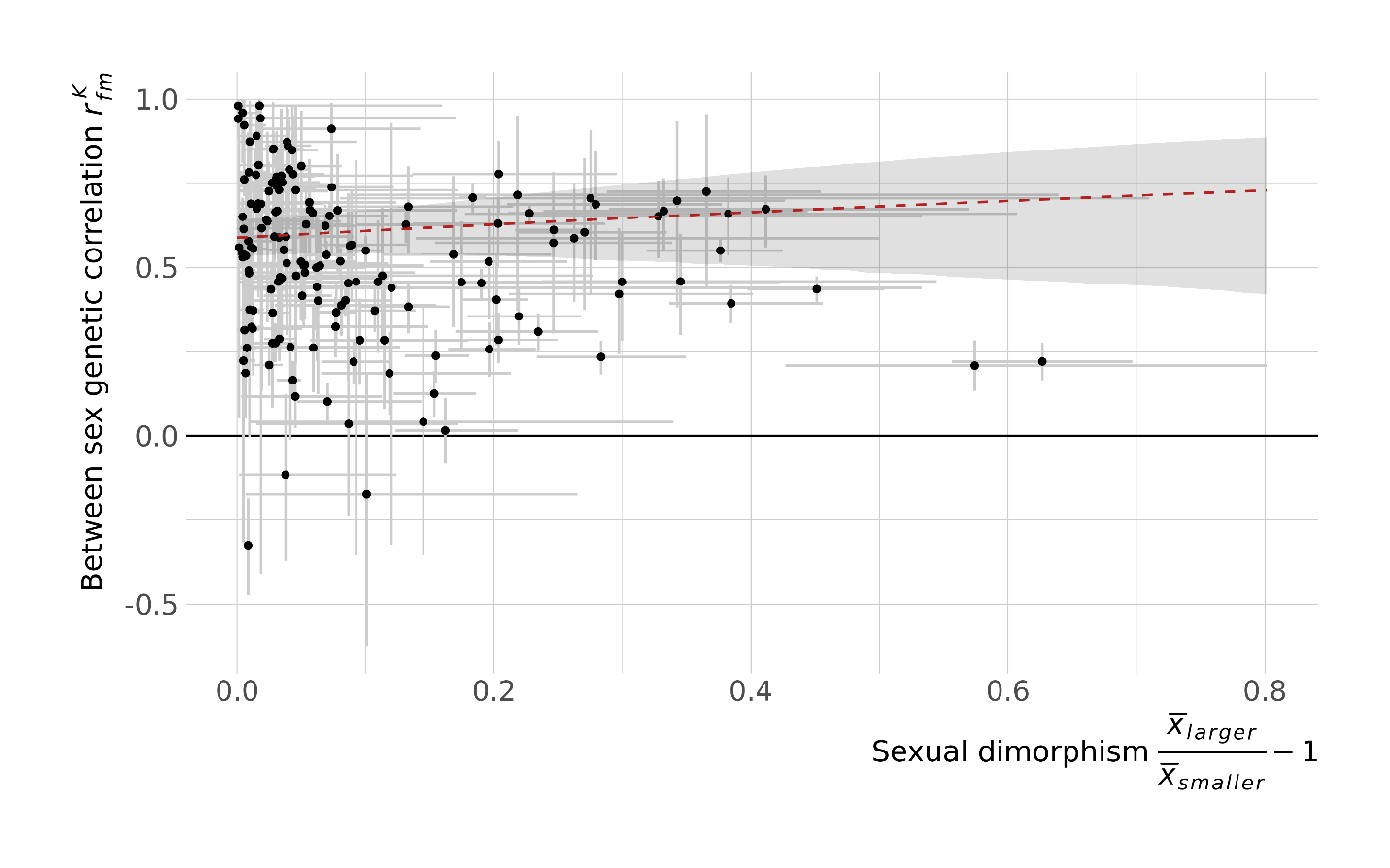

**Figure S6:** As Figure 2, but without accounting for body mass. The between sexual genetic correlation does not depend on sexual dimorphism in the trait. Each point is a trait, with error bars indicating the 95% credible interval (CI) in the estimates. The line represents the model fit of a linear model on the Fisher-transformed $r_{fm}^{K}$, with the shaded region indicating the 95% credible interval, including propagation of trait level uncertainty.

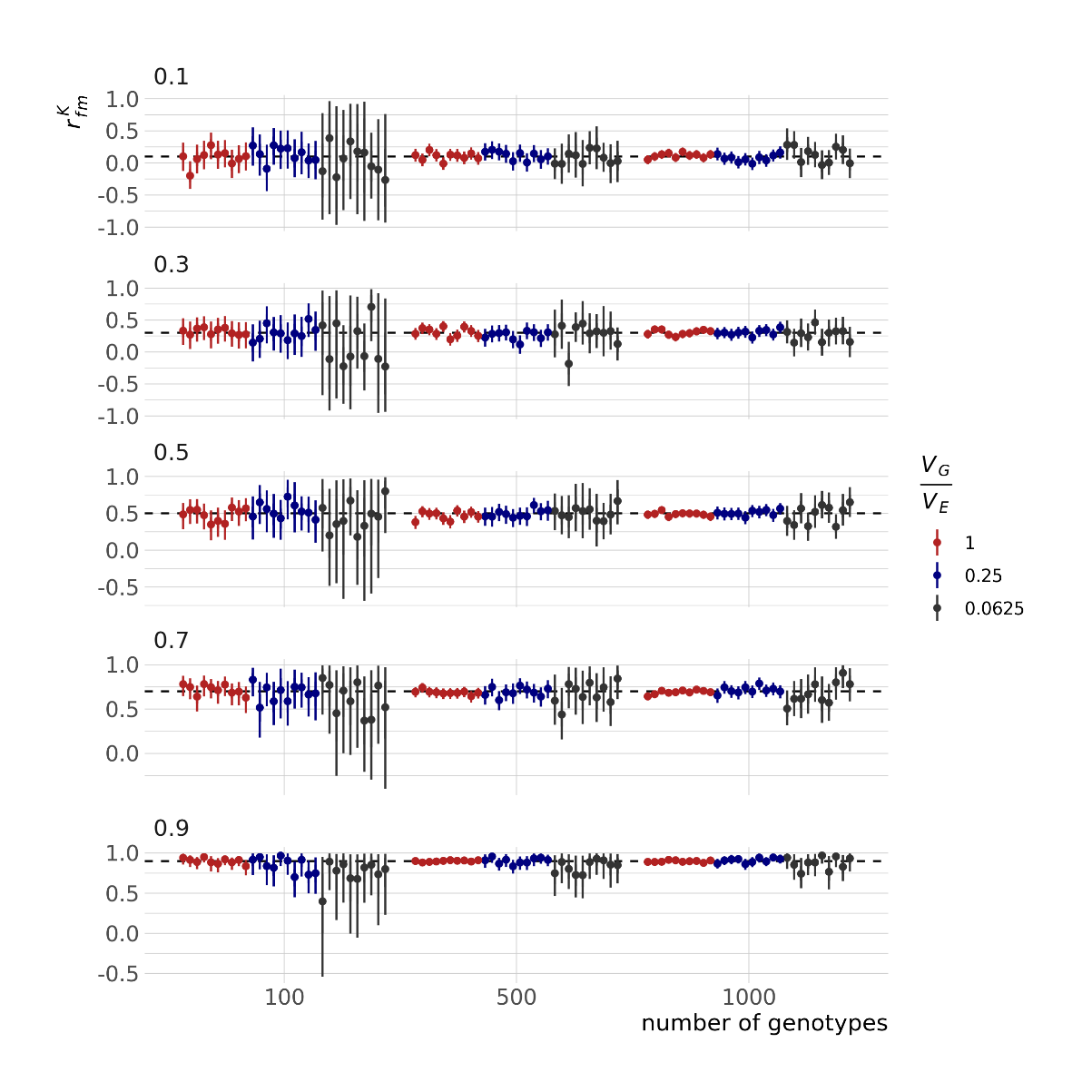

**Figirue S7:** A simulation study on the potential for biased estimates for $r_{fm}^{K}$ from our modelling approach. We simulated data, with varying number of genotypes available (100, 500 or 1000), varying true values for $r_{fm}^{K}$ (0.1, 0.3, 0.5, 0.7, 0.9), and varying proportions of genetic variance (compared to non-genetic variance, 1:1, 1:4, 1:16). We replicated the limited data available for each genotype in the IMPC data, and simulated 7 females and 7 males per genotype. The figure shows the point estimate and 95% Credible Interval for 10 fitted models for each scenario.

**Table S1**: Estimated parameters for each trait. *n* denotes the total sample size (number of individuals tested), *genotypes* the number of different genotypes, *SD* is sexual dimorphism, *rFM* is the between sex genetic correlation. *SD* and *rFM* are given as the estimate with 95% Credible Intervals.

| Trait ID | Trait name | Trait category | n | genotypes | SD index | rFM |
| --- | --- | --- | --- | --- | --- | --- |
| IMPC_ACS_001_001 | Response amplitude - BN | physiology | 68729 | 3611 |  | 0.748 [0.673, 0.829] |
| IMPC_ACS_002_001 | Response amplitude - PP1 | physiology | 68731 | 3611 |  | 0.765 [0.704, 0.827] |
| IMPC_ACS_003_001 | Response amplitude - PP2 | physiology | 68725 | 3611 |  | 0.747 [0.679, 0.811] |
| IMPC_ACS_004_001 | Response amplitude - PP3 | physiology | 68731 | 3611 |  | 0.749 [0.698, 0.804] |
| IMPC_ACS_005_001 | Response amplitude - PP4 | physiology | 31535 | 1302 |  | 0.882 [0.812, 0.947] |
| IMPC_ACS_006_001 | Response amplitude - S | physiology | 68343 | 3594 |  | 0.872 [0.842, 0.9] |
| IMPC_ACS_007_001 | Response amplitude - PP1_S | physiology | 68350 | 3594 |  | 0.879 [0.852, 0.907] |
| IMPC_ACS_008_001 | Response amplitude - PP2_S | physiology | 68344 | 3594 |  | 0.89 [0.862, 0.917] |
| IMPC_ACS_009_001 | Response amplitude - PP3_S | physiology | 68350 | 3594 |  | 0.871 [0.842, 0.898] |
| IMPC_ACS_010_001 | Response amplitude - PP4_S | physiology | 31535 | 1302 |  | 0.856 [0.809, 0.906] |
| IMPC_ACS_033_001 | % Pre-pulse inhibition - PPI1 | physiology | 68333 | 3594 | 0.044 [0, 0.107] | 0.786 [0.635, 0.944] |
| IMPC_ACS_034_001 | % Pre-pulse inhibition - PPI2 | physiology | 68327 | 3594 | 0.022 [0, 0.042] | 0.817 [0.757, 0.876] |
| IMPC_ACS_035_001 | % Pre-pulse inhibition - PPI3 | physiology | 68333 | 3594 | 0.01 [0, 0.023] | 0.766 [0.715, 0.818] |
| IMPC_ACS_036_001 | % Pre-pulse inhibition - PPI4 | physiology | 31536 | 1302 | 0.02 [0, 0.039] | 0.752 [0.673, 0.835] |
| IMPC_ACS_037_001 | % Pre-pulse inhibition - Global | physiology | 31530 | 1301 | 0.016 [0, 0.046] | 0.724 [0.626, 0.81] |
| IMPC_ABR_002_001 | Click-evoked ABR threshold | physiology | 17535 | 3090 |  | 0.919 [0.891, 0.945] |
| IMPC_ABR_004_001 | 6kHz-evoked ABR Threshold | physiology | 30798 | 4404 |  | 0.904 [0.869, 0.937] |
| IMPC_ABR_006_001 | 12kHz-evoked ABR Threshold | physiology | 30806 | 4404 |  | 0.923 [0.893, 0.952] |
| IMPC_ABR_008_001 | 18kHz-evoked ABR Threshold | physiology | 30781 | 4403 |  | 0.899 [0.869, 0.931] |
| IMPC_ABR_010_001 | 24kHz-evoked ABR Threshold | physiology | 30685 | 4400 |  | 0.879 [0.841, 0.917] |
| IMPC_ABR_012_001 | 30kHz-evoked ABR Threshold | physiology | 30185 | 4397 |  | 0.833 [0.785, 0.885] |
| IMPC_DXA_002_001 | Fat mass | morphology | 81898 | 4346 | 0.155 [0.123, 0.198] | 0.313 [0.267, 0.359] |
| IMPC_DXA_005_001 | Bone Mineral Content (excluding skull) | morphology | 81045 | 4316 | 0.069 [0.059, 0.081] | 0.305 [0.265, 0.348] |
| IMPC_DXA_006_001 | Body length | morphology | 67662 | 3724 | 0.002 [0, 0.004] | 0.499 [0.458, 0.538] |
| IMPC_DXA_010_001 | Bone Area | morphology | 81046 | 4316 | 0.034 [0.024, 0.042] | 0.437 [0.387, 0.486] |
| IMPC_CBC_001_001 | Sodium | physiology | 49468 | 2470 | 0.009 [0.005, 0.012] | -0.098 [-0.202, 0.016] |
| IMPC_CBC_002_001 | Potassium | physiology | 49249 | 2470 | 0.093 [0.076, 0.112] | 0.412 [0.318, 0.512] |
| IMPC_CBC_004_001 | Urea (Blood Urea Nitrogen - BUN) | physiology | 73486 | 2949 | 0.01 [0, 0.028] | 0.461 [0.388, 0.53] |
| IMPC_CBC_005_001 | Creatinine | physiology | 65688 | 2523 | 0.107 [0.078, 0.134] | 0.284 [0.207, 0.366] |
| IMPC_CBC_006_001 | Total protein | physiology | 73404 | 2945 | 0.005 [0, 0.009] | 0.372 [0.289, 0.456] |
| IMPC_CBC_007_001 | Albumin | physiology | 74225 | 2950 | 0.064 [0.058, 0.069] | 0.301 [0.22, 0.378] |
| IMPC_CBC_008_001 | Total bilirubin | physiology | 72729 | 2939 | 0.011 [0, 0.028] | 0.526 [0.431, 0.623] |
| IMPC_CBC_009_001 | Calcium | physiology | 73880 | 2948 | 0.001 [0, 0.003] | 0.327 [0.211, 0.43] |
| IMPC_CBC_010_001 | Phosphorus | physiology | 73250 | 2947 | 0.027 [0.015, 0.041] | 0.499 [0.417, 0.58] |
| IMPC_CBC_011_001 | Iron | physiology | 47260 | 2301 | 0.185 [0.163, 0.207] | 0.162 [0.078, 0.248] |
| IMPC_CBC_012_001 | Aspartate aminotransferase | physiology | 73624 | 2950 | 0.007 [0, 0.024] | 0.574 [0.483, 0.669] |
| IMPC_CBC_013_001 | Alanine aminotransferase | physiology | 74021 | 2950 | 0.127 [0.091, 0.161] | 0.519 [0.427, 0.613] |
| IMPC_CBC_014_001 | Alkaline phosphatase | physiology | 73703 | 2937 | 0.39 [0.353, 0.425] | 0.593 [0.549, 0.635] |
| IMPC_CBC_015_001 | Total cholesterol | physiology | 73572 | 2947 | 0.058 [0.044, 0.073] | 0.643 [0.599, 0.692] |
| IMPC_CBC_016_001 | HDL-cholesterol | physiology | 67324 | 2793 | 0.127 [0.101, 0.159] | 0.614 [0.564, 0.664] |
| IMPC_CBC_017_001 | Triglycerides | physiology | 72229 | 2933 | 0.136 [0.085, 0.193] | 0.366 [0.297, 0.429] |
| IMPC_CBC_018_001 | Glucose | physiology | 73210 | 2949 | 0.044 [0, 0.088] | 0.135 [0.067, 0.206] |
| IMPC_CBC_020_001 | Fructosamine | physiology | 31243 | 1655 | 0.049 [0.031, 0.062] | 0.35 [0.268, 0.428] |
| IMPC_CBC_021_001 | Lipase | physiology | 6620 | 264 | 0.014 [0, 0.031] | 0.886 [0.737, 0.991] |
| IMPC_CBC_022_001 | Lactate dehydrogenase | physiology | 3675 | 115 | 0.123 [0.019, 0.241] | 0.466 [-0.238, 0.955] |
| IMPC_CBC_023_001 | Alpha-amylase | physiology | 37847 | 1790 | 0.144 [0.121, 0.167] | 0.643 [0.583, 0.706] |
| IMPC_CBC_024_001 | UIBC (unsaturated iron binding capacity) | physiology | 6681 | 264 | 0.07 [0.05, 0.091] | 0.429 [0.217, 0.632] |
| IMPC_CBC_025_001 | LDL-cholesterol | physiology | 21025 | 1079 | 0.113 [0.047, 0.182] | 0.361 [0.27, 0.448] |
| IMPC_CBC_026_001 | Free fatty acids | physiology | 23343 | 1416 | 0.051 [0.008, 0.088] | 0.311 [0.192, 0.428] |
| IMPC_CBC_028_001 | Creatine kinase | physiology | 31464 | 1616 | 0.052 [0, 0.102] | 0.858 [0.585, 1] |
| IMPC_CSD_032_001 | Locomotor activity | behavior | 70051 | 3822 | 0.139 [0.108, 0.173] | 0.333 [0.267, 0.397] |
| IMPC_ECH_001_001 | End-Systolic Diameter | morphology | 5954 | 733 | 0.014 [0, 0.031] | 0.625 [0.441, 0.796] |
| IMPC_ECH_002_001 | End-Diastolic Diameter | morphology | 5954 | 733 | 0.011 [0, 0.021] | 0.634 [0.45, 0.812] |
| IMPC_ECH_003_001 | Stroke Volume | physiology | 13123 | 1014 | 0.022 [0, 0.047] | 0.538 [0.337, 0.73] |
| IMPC_ECH_004_001 | Ejection Fraction | physiology | 15275 | 1186 |  | 0.636 [0.49, 0.783] |
| IMPC_ECH_005_001 | Fractional Shortening | physiology | 19414 | 1193 |  | 0.687 [0.54, 0.832] |
| IMPC_ECH_006_001 | Cardiac Output | physiology | 13101 | 612 | 0.022 [0, 0.048] | 0.527 [0.026, 0.979] |
| IMPC_ECH_008_001 | LVIDd | physiology | 19414 | 1193 | 0.011 [0, 0.024] | 0.544 [0.408, 0.672] |
| IMPC_ECH_009_001 | LVPWd | physiology | 19415 | 1193 | 0.004 [0, 0.013] | 0.317 [0.095, 0.534] |
| IMPC_ECH_010_001 | LVAWs | physiology | 8078 | 905 | 0.014 [0, 0.033] | 0.585 [0.369, 0.8] |
| IMPC_ECH_011_001 | LVIDs | physiology | 19415 | 1193 | 0.013 [0, 0.036] | 0.605 [0.461, 0.731] |
| IMPC_ECH_012_001 | LVPWs | physiology | 15274 | 1186 | 0.009 [0, 0.021] | 0.411 [0.234, 0.584] |
| IMPC_ECH_013_001 | HR | physiology | 19406 | 789 | 0.011 [0, 0.024] | 0.509 [0.283, 0.732] |
| IMPC_ECH_014_001 | Body Temp | physiology | 3798 | 194 |  | 0.966 [0.792, 1] |
| IMPC_ECH_018_001 | Respiration Rate | physiology | 10723 | 589 | 0.031 [0, 0.075] | 0.689 [0.4, 0.935] |
| IMPC_ECG_001_001 | Number of signals | physiology | 58260 | 3272 | 0.011 [0, 0.026] | 0.451 [0.282, 0.629] |
| IMPC_ECG_002_001 | HR | physiology | 61035 | 3387 | 0.015 [0.011, 0.02] | 0.486 [0.406, 0.569] |
| IMPC_ECG_003_001 | CV | physiology | 37965 | 2288 |  | 0.5 [0.279, 0.719] |
| IMPC_ECG_004_001 | RR | physiology | 61034 | 3387 | 0.015 [0.011, 0.021] | 0.475 [0.392, 0.561] |
| IMPC_ECG_005_001 | PQ | physiology | 37178 | 2247 | 0.019 [0.005, 0.032] | 0.605 [0.395, 0.8] |
| IMPC_ECG_006_001 | PR | physiology | 61036 | 3387 | 0.015 [0.009, 0.02] | 0.597 [0.363, 0.843] |
| IMPC_ECG_007_001 | QRS | physiology | 61022 | 3387 | 0.007 [0.001, 0.013] | 0.624 [0.347, 0.913] |
| IMPC_ECG_008_001 | ST | physiology | 56467 | 3204 | 0.014 [0.01, 0.019] | 0.514 [0.3, 0.714] |
| IMPC_ECG_009_002 | QTc | physiology | 3889 | 621 | 0.005 [0.001, 0.008] | 0.635 [-0.021, 0.999] |
| IMPC_ECG_010_001 | HRV | physiology | 37178 | 2247 | 0.311 [0.21, 0.424] | 0.463 [0.23, 0.685] |
| IMPC_ECG_011_001 | QTc Dispersion | physiology | 38928 | 2315 | 0.023 [0, 0.044] | 0.694 [0.396, 0.992] |
| IMPC_ECG_012_001 | Mean SR amplitude | physiology | 37172 | 2247 | 0.134 [0.071, 0.215] | 0.405 [0.308, 0.504] |
| IMPC_ECG_013_001 | Mean R amplitude | physiology | 40058 | 2315 | 0.156 [0.076, 0.243] | 0.328 [0.224, 0.428] |
| IMPC_ECG_014_001 | rMSSD | physiology | 37176 | 2247 | 0.193 [0.12, 0.276] | 0.648 [0.403, 0.908] |
| IMPC_EYE_054_001 | Min left eye lens density | morphology | 4271 | 149 |  | 0.898 [0.774, 0.999] |
| IMPC_EYE_055_001 | Max left eye lens density | morphology | 4271 | 149 |  | 0.92 [0.785, 1] |
| IMPC_EYE_056_001 | Mean left eye lens density | morphology | 4271 | 149 |  | 0.937 [0.833, 1] |
| IMPC_EYE_057_001 | Min right eye lens density | morphology | 4242 | 149 |  | 0.883 [0.751, 0.998] |
| IMPC_EYE_058_001 | Max right eye lens density | morphology | 4242 | 149 |  | 0.719 [0.536, 0.866] |
| IMPC_EYE_059_001 | Mean right eye lens density | morphology | 4242 | 149 |  | 0.798 [0.674, 0.911] |
| IMPC_EYE_063_001 | Right inner nuclear layer | morphology | 3995 | 121 | 0.011 [0, 0.035] | 0.866 [0.608, 0.998] |
| IMPC_EYE_064_001 | Right outer nuclear layer | morphology | 3997 | 121 | 0.015 [0, 0.067] | 0.411 [-0.774, 1] |
| IMPC_EYE_068_001 | Left total retinal thickness | morphology | 8765 | 305 | 0.001 [0, 0.003] | 0.947 [0.852, 1] |
| IMPC_EYE_069_001 | Left inner nuclear layer | morphology | 4073 | 121 | 0.022 [0, 0.049] | 0.909 [0.66, 1] |
| IMPC_EYE_070_001 | Left outer nuclear layer | morphology | 4072 | 121 | 0.018 [0, 0.08] | -0.04 [-0.925, 0.96] |
| IMPC_FEA_001_001 | Conditioning Baseline Freeze Count | behavior | 1551 | 135 | 0.196 [0, 0.702] | 0.343 [-0.339, 0.95] |
| IMPC_FEA_006_001 | Conditioning Baseline Maximum Motion Index | behavior | 1551 | 108 |  | 0.655 [0.267, 0.999] |
| IMPC_FEA_007_001 | Context Freeze Count | behavior | 1551 | 135 | 0.027 [0, 0.09] | 0.866 [0.534, 1] |
| IMPC_FEA_008_001 | Context Freezing Time | behavior | 1551 | 135 | 0.034 [0, 0.12] | 0.929 [0.731, 1] |
| IMPC_FEA_012_001 | Context Maximum Motion Index | behavior | 1551 | 108 |  | 0.626 [-0.403, 1] |
| IMPC_FEA_018_001 | Cue Baseline Maximum Motion Index | behavior | 1549 | 108 |  | 0.683 [0.243, 0.998] |
| IMPC_FEA_021_001 | Cue Tone % Freezing Time | behavior | 1550 | 108 |  | 0.72 [0.39, 1] |
| IMPC_FEA_094_001 | Conditioning Tone Maximum Motion Index | behavior | 1551 | 108 |  | 0.487 [-0.282, 0.999] |
| IMPC_FEA_095_001 | Conditioning Shock Average Motion Index | behavior | 1551 | 108 |  | 0.586 [0.231, 0.995] |
| IMPC_FEA_096_001 | Conditioning Shock Minimum Motion Index | behavior | 1551 | 108 |  | 0.21 [-0.852, 1] |
| IMPC_FEA_097_001 | Conditioning Shock Maximum Motion Index | behavior | 1551 | 108 |  | 0.719 [0.218, 1] |
| IMPC_FEA_099_001 | Conditioning Post-shock Freezing Time | behavior | 1551 | 135 | 0.09 [0, 0.278] | 0.88 [0.65, 1] |
| IMPC_FEA_103_001 | Conditioning Post-shock Maximum Motion Index | behavior | 1551 | 108 |  | 0.535 [-0.586, 1] |
| IMPC_GRS_008_001 | Forelimb grip strength measurement mean | physiology | 83155 | 4157 | 0.015 [0.005, 0.026] | 0.499 [0.441, 0.551] |
| IMPC_GRS_009_001 | Forelimb and hindlimb grip strength measurement mean | physiology | 83067 | 4158 | 0.038 [0.029, 0.046] | 0.552 [0.507, 0.598] |
| IMPC_GRS_011_001 | Forelimb and hindlimb grip strength normalised against body weight | physiology | 82998 | 4156 |  | 0.461 [0.416, 0.505] |
| IMPC_HWT_008_001 | Heart weight | morphology | 71786 | 3867 | 0.073 [0.063, 0.082] | 0.655 [0.596, 0.713] |
| IMPC_HEM_001_001 | White blood cell count | physiology | 73628 | 3652 | 0.24 [0.173, 0.294] | 0.344 [0.282, 0.4] |
| IMPC_HEM_002_001 | Red blood cell count | physiology | 75119 | 3683 | 0.024 [0.011, 0.035] | 0.196 [0.124, 0.259] |
| IMPC_HEM_003_001 | Hemoglobin | physiology | 75074 | 3683 | 0.004 [0, 0.01] | 0.228 [0.164, 0.288] |
| IMPC_HEM_004_001 | Hematocrit | physiology | 75103 | 3683 |  | 0.402 [0.343, 0.463] |
| IMPC_HEM_008_001 | Platelet count | physiology | 74953 | 3681 | 0.207 [0.15, 0.253] | 0.319 [0.237, 0.397] |
| IMPC_HEM_029_001 | Neutrophil differential count | physiology | 37519 | 1718 |  | 0.481 [0.41, 0.553] |
| IMPC_HEM_030_001 | Neutrophil cell count | physiology | 34421 | 1495 | 0.206 [0.165, 0.26] | 0.386 [0.291, 0.472] |
| IMPC_HEM_031_001 | Lymphocyte differential count | physiology | 38837 | 1763 |  | 0.473 [0.402, 0.545] |
| IMPC_HEM_032_001 | Lymphocyte cell count | physiology | 34256 | 1483 | 0.183 [0.147, 0.222] | 0.287 [0.205, 0.371] |
| IMPC_HEM_033_001 | Monocyte differential count | physiology | 38805 | 1761 |  | 0.405 [0.312, 0.5] |
| IMPC_HEM_034_001 | Monocyte cell count | physiology | 34262 | 1483 | 0.132 [0.081, 0.18] | 0.045 [-0.06, 0.144] |
| IMPC_HEM_035_001 | Eosinophil differential count | physiology | 37821 | 1760 |  | 0.472 [0.342, 0.587] |
| IMPC_HEM_036_001 | Eosinophil cell count | physiology | 34246 | 1483 | 0.145 [0.07, 0.246] | 0.236 [0.108, 0.366] |
| IMPC_HEM_037_001 | Basophil cell count | physiology | 33765 | 1491 | 0.082 [0, 0.204] | 0.36 [0.217, 0.484] |
| IMPC_HEM_038_001 | Basophil differential count | physiology | 37050 | 1764 |  | 0.512 [0.394, 0.63] |
| IMPC_HEM_039_001 | Large Unstained Cell (LUC) count | physiology | 17501 | 703 | 0.278 [0.167, 0.45] | 0.68 [0.564, 0.792] |
| IMPC_HEM_040_001 | Large Unstained Cell (LUC) differential count | physiology | 17498 | 702 |  | 0.655 [0.505, 0.793] |
| IMPC_IMM_001_001 | Spleen weight | morphology | 6040 | 616 | 0.347 [0.306, 0.387] | 0.398 [0.227, 0.558] |
| IMPC_IMM_002_001 | Percentage of live gated events in Panel A | immunology | 4140 | 565 |  | 0.304 [-0.689, 0.988] |
| IMPC_IMM_003_001 | T cells (panel A) | immunology | 4422 | 549 | 0.013 [0, 0.039] | 0.823 [0.532, 1] |
| IMPC_IMM_004_001 | NKT cells (panel A) | immunology | 4421 | 578 | 0.15 [0.057, 0.248] | 0.648 [0.511, 0.774] |
| IMPC_IMM_006_001 | Others | immunology | 4270 | 578 | 0.042 [0.003, 0.079] | 0.741 [0.457, 0.959] |
| IMPC_IMM_007_001 | CD4 T cells | immunology | 4422 | 549 | 0.016 [0, 0.045] | 0.742 [0.45, 0.986] |
| IMPC_IMM_008_001 | CD8 T cells | immunology | 4417 | 549 | 0.018 [0, 0.048] | 0.859 [0.653, 1] |
| IMPC_IMM_009_001 | DN T cells | immunology | 4399 | 546 | 0.281 [0.199, 0.363] | 0.694 [0.465, 0.916] |
| IMPC_IMM_010_001 | DP T cells | immunology | 3194 | 521 | 0.023 [0, 0.072] | 0.617 [0.321, 0.903] |
| IMPC_IMM_011_001 | CD4 NKT cells | immunology | 4269 | 549 | 0.314 [0.172, 0.466] | 0.664 [0.543, 0.779] |
| IMPC_IMM_012_001 | CD8 NKT cells | immunology | 4251 | 546 | 0.062 [0, 0.15] | 0.631 [0.426, 0.839] |
| IMPC_IMM_013_001 | DN NKT cells | immunology | 4251 | 574 | 0.102 [0.018, 0.191] | 0.777 [0.631, 0.933] |
| IMPC_IMM_014_001 | CD4 CD25+ T cells | immunology | 4414 | 549 | 0.052 [0, 0.1] | -0.142 [-0.543, 0.24] |
| IMPC_IMM_015_001 | CD4 CD25- T cells | immunology | 4414 | 549 | 0.016 [0, 0.049] | 0.764 [0.445, 1] |
| IMPC_IMM_016_001 | CD8 CD25+ T cells | immunology | 3733 | 470 | 0.112 [0.031, 0.208] | 0.905 [0.612, 1] |
| IMPC_IMM_022_001 | CD8 CD25+ NKT cells | immunology | 3715 | 467 | 0.117 [0.008, 0.22] | 0.793 [0.234, 1] |
| IMPC_IMM_023_001 | CD8 CD25- NKT cells | immunology | 3715 | 467 | 0.032 [0, 0.095] | 0.592 [0.39, 0.775] |
| IMPC_IMM_025_001 | DN CD25- NKT cells | immunology | 3715 | 466 | 0.192 [0.094, 0.321] | 0.728 [0.528, 0.915] |
| IMPC_IMM_026_001 | Total number of acquired events in Panel A | immunology | 4189 | 577 | 0.013 [0, 0.038] | 0.38 [0.03, 0.698] |
| IMPC_IMM_027_001 | Total number of acquired events in Panel B | immunology | 4198 | 577 | 0.024 [0, 0.064] | 0.794 [0.683, 0.901] |
| IMPC_IMM_028_001 | CD4 CD44+CD62L- T cells | immunology | 4414 | 549 | 0.082 [0.005, 0.147] | 0.715 [0.491, 0.898] |
| IMPC_IMM_029_001 | CD4 CD44+CD62L+ T cells | immunology | 4174 | 533 | 0.053 [0, 0.111] | 0.753 [0.521, 0.967] |
| IMPC_IMM_030_001 | CD4 CD44-CD62L+ T cells | immunology | 3886 | 470 | 0.034 [0, 0.074] | 0.836 [0.457, 1] |
| IMPC_IMM_031_001 | CD4 CD44-CD62L- T cells | immunology | 2491 | 432 | 0.029 [0, 0.083] | 0.66 [-0.076, 1] |
| IMPC_IMM_032_001 | CD8 CD44+CD62L- T cells | immunology | 4414 | 549 | 0.029 [0, 0.082] | 0.74 [0.55, 0.92] |
| IMPC_IMM_033_001 | CD8 CD44+CD62L+ T cells | immunology | 4414 | 549 | 0.045 [0, 0.104] | 0.751 [0.598, 0.89] |
| IMPC_IMM_034_001 | CD8 CD44-CD62L+ T cells | immunology | 4414 | 549 | 0.019 [0, 0.058] | 0.845 [0.66, 1] |
| IMPC_IMM_035_001 | CD8 CD44-CD62L- T cells | immunology | 3192 | 521 | 0.051 [0, 0.126] | 0.609 [0.395, 0.796] |
| IMPC_IMM_036_001 | DN CD44+CD62L- T cells | immunology | 4243 | 545 | 0.377 [0.261, 0.508] | 0.741 [0.559, 0.911] |
| IMPC_IMM_038_001 | DN CD44-CD62L+ T cells | immunology | 4243 | 545 | 0.315 [0.233, 0.402] | 0.711 [0.421, 0.96] |
| IMPC_IMM_039_001 | DN CD44-CD62L- T cells | immunology | 3173 | 517 | 0.405 [0.312, 0.496] | 0.797 [0.543, 0.995] |
| IMPC_IMM_040_001 | CD4 CD44+CD62L- NKT cells | immunology | 4249 | 547 | 0.365 [0.187, 0.55] | 0.678 [0.557, 0.794] |
| IMPC_IMM_041_001 | CD4 CD44+CD62L+ NKT cells | immunology | 4261 | 549 | 0.04 [0, 0.111] | 0.385 [0.15, 0.592] |
| IMPC_IMM_042_001 | CD4 CD44-CD62L+ NKT cells | immunology | 4261 | 549 | 0.122 [0, 0.317] | -0.115 [-0.507, 0.315] |
| IMPC_IMM_043_001 | CD8 CD44+CD62L- NKT cells | immunology | 4243 | 546 | 0.038 [0, 0.112] | 0.517 [0.205, 0.857] |
| IMPC_IMM_044_001 | CD8 CD44+CD62L+ NKT cells | immunology | 4243 | 546 | 0.096 [0, 0.197] | 0.596 [0.426, 0.773] |
| IMPC_IMM_045_001 | CD8 CD44-CD62L+ NKT cells | immunology | 4243 | 546 | 0.065 [0, 0.192] | 0.257 [-0.16, 0.641] |
| IMPC_IMM_047_001 | DN CD44+CD62L+ NKT cells | immunology | 4243 | 545 | 0.028 [0, 0.083] | 0.763 [0.62, 0.9] |
| IMPC_IMM_048_001 | DN CD44-CD62L+ NKT cells | immunology | 4243 | 545 | 0.644 [0.328, 1.087] | 0.155 [-0.798, 0.911] |
| IMPC_IMM_050_001 | Neutrophils | immunology | 4265 | 449 | 0.047 [0, 0.155] | 0.433 [0.2, 0.647] |
| IMPC_IMM_051_001 | Monocytes | immunology | 4245 | 448 | 0.046 [0, 0.11] | 0.29 [0.068, 0.501] |
| IMPC_IMM_052_001 | Eosinophils | immunology | 4265 | 449 | 0.083 [0.007, 0.153] | 0.366 [0.063, 0.656] |
| IMPC_IMM_053_001 | NK Cells (panel B) | immunology | 3872 | 505 | 0.023 [0, 0.067] | 0.62 [0.461, 0.768] |
| IMPC_IMM_054_001 | NK Subsets (Q1) | immunology | 4169 | 510 | 0.105 [0.019, 0.194] | -0.094 [-0.309, 0.111] |
| IMPC_IMM_055_001 | NK Subsets (Q2) | immunology | 4169 | 510 | 0.041 [0, 0.116] | 0.664 [0.504, 0.829] |
| IMPC_IMM_056_001 | NK Subsets (Q3) | immunology | 3649 | 409 | 0.212 [0.097, 0.364] | 0.481 [0.27, 0.669] |
| IMPC_IMM_057_001 | NK Subsets (Q4) | immunology | 3649 | 409 | 0.056 [0, 0.128] | 0.586 [0.426, 0.734] |
| IMPC_IMM_058_001 | NKT Cells (panel B) | immunology | 3872 | 502 | 0.191 [0.105, 0.285] | 0.843 [0.725, 0.955] |
| IMPC_IMM_059_001 | NKT Subsets (Q1) | immunology | 4169 | 510 | 0.091 [0.027, 0.161] | 0.751 [0.599, 0.889] |
| IMPC_IMM_060_001 | NKT Subsets (Q3) | immunology | 3646 | 407 | 0.353 [0.223, 0.523] | 0.571 [0.407, 0.731] |
| IMPC_IMM_061_001 | T Cells (panel B) | immunology | 4265 | 576 | 0.013 [0, 0.039] | 0.761 [0.628, 0.884] |
| IMPC_IMM_066_001 | Follicular B Cells | immunology | 2561 | 460 | 0.029 [0, 0.068] | 0.886 [0.741, 0.992] |
| IMPC_IMM_068_001 | Transitional B Cells | immunology | 1601 | 317 | 0.137 [0, 0.318] | 0.058 [-0.616, 0.583] |
| IMPC_IMM_070_001 | MZB | immunology | 1165 | 221 | 0.036 [0, 0.117] | 0.836 [0.614, 1] |
| IMPC_IMM_071_001 | MZB (CD21/35 high) | immunology | 2200 | 224 | 0.061 [0, 0.174] | 0.858 [0.594, 1] |
| IMPC_IMM_072_001 | cDCs | immunology | 4260 | 577 | 0.194 [0.12, 0.278] | 0.529 [0.33, 0.71] |
| IMPC_IMM_073_001 | cDCs CD11b Type | immunology | 4260 | 577 | 0.295 [0.195, 0.405] | 0.524 [0.328, 0.702] |
| IMPC_IMM_074_001 | pDCs | immunology | 2569 | 439 | 0.124 [0.051, 0.207] | 0.9 [0.746, 0.991] |
| IMPC_IMM_075_001 | RP Macrophage (F4/80+) | physiology | 951 | 146 | 0.057 [0, 0.114] | 0.637 [0.33, 0.897] |
| IMPC_CAL_008_001 | Total food intake | behavior | 16108 | 876 | 0.027 [0, 0.065] | 0.728 [0.631, 0.822] |
| IMPC_CAL_017_001 | Respiratory Exchange Ratio | physiology | 29638 | 2179 |  | 0.611 [0.501, 0.703] |
| IMPC_CAL_021_001 | Total water intake | behavior | 9823 | 743 | 0.037 [0, 0.09] | 0.491 [0.272, 0.697] |
| IMPC_IPG_010_001 | Fasted blood glucose concentration | physiology | 81056 | 4424 | 0.035 [0.02, 0.05] | 0.399 [0.353, 0.446] |
| IMPC_IPG_011_001 | Initial response to glucose challenge | physiology | 80897 | 4424 | 0.173 [0.142, 0.206] | 0.125 [0.058, 0.195] |
| IMPC_IPG_012_001 | Area under glucose response curve | physiology | 80709 | 4424 | 0.462 [0.406, 0.522] | 0.224 [0.165, 0.278] |
| IMPC_OFD_009_001 | Whole arena average speed | behavior | 70202 | 3367 | 0.072 [0.058, 0.086] | 0.52 [0.475, 0.559] |
| IMPC_OFD_010_001 | Periphery distance travelled | behavior | 69000 | 3332 | 0.09 [0.077, 0.105] | 0.549 [0.503, 0.593] |
| IMPC_OFD_011_001 | Periphery resting time | behavior | 37620 | 1577 | 0.033 [0, 0.055] | 0.66 [0.601, 0.713] |
| IMPC_OFD_012_001 | Periphery permanence time | behavior | 70171 | 3367 | 0.002 [0, 0.005] | 0.644 [0.585, 0.707] |
| IMPC_OFD_014_001 | Center distance travelled | behavior | 68997 | 3332 | 0.054 [0.011, 0.097] | 0.652 [0.606, 0.699] |
| IMPC_OFD_015_001 | Center resting time | behavior | 37617 | 1577 | 0.034 [0, 0.102] | 0.46 [0.353, 0.557] |
| IMPC_OFD_016_001 | Center permanence time | behavior | 70168 | 3367 | 0.013 [0, 0.042] | 0.633 [0.575, 0.697] |
| IMPC_OFD_017_001 | Center average speed | behavior | 64009 | 3088 | 0.074 [0.055, 0.096] | 0.629 [0.553, 0.7] |
| IMPC_OFD_018_001 | Latency to center entry | behavior | 38012 | 1591 | 0.066 [0, 0.203] | 0.441 [0.228, 0.648] |
| IMPC_OFD_019_001 | Number of center entries | behavior | 37999 | 1590 | 0.016 [0, 0.06] | 0.674 [0.613, 0.737] |
| IMPC_OFD_020_001 | Distance travelled - total | behavior | 68018 | 3285 | 0.074 [0.062, 0.088] | 0.566 [0.522, 0.606] |
| IMPC_OFD_021_001 | Number of rears - total | behavior | 49728 | 2537 | 0.04 [0, 0.081] | 0.649 [0.591, 0.701] |
| IMPC_OFD_022_001 | Percentage center time | behavior | 65188 | 3123 |  | 0.633 [0.57, 0.692] |

**Table S2:** List of genotypes with consistently low or consistently high discordant ranks. *Traits* column refers to number of traits that genotype was tested for. *Lower CI* and *upper CI* denote the lower and upper bounds of the Credible Interval associated with the mean discordant rank.

| background | gene | allele | zygosity | traits | mean discordant rank | lower CI | upper CI |
| --- | --- | --- | --- | --- | --- | --- | --- |
| involves: C57BL/6N | wildtype | wildtype | homozygote | 174 | 0.2634 | 0.2432 | 0.2833 |
| involves: C57BL/6NTac | wildtype | wildtype | homozygote | 186 | 0.2977 | 0.2826 | 0.3141 |
| involves: C57BL/6NJ | wildtype | wildtype | homozygote | 148 | 0.3375 | 0.3105 | 0.3662 |
| involves: C57BL/6NCrl | wildtype | wildtype | homozygote | 192 | 0.3560 | 0.3336 | 0.3776 |
| involves: C57BL/6N;C57BL/6NTac | wildtype | wildtype | homozygote | 170 | 0.4501 | 0.4202 | 0.4784 |
| involves: C57BL/6N | MGI:1338891 | MGI:5793070 | homozygote | 116 | 0.5479 | 0.5019 | 0.5940 |
| involves: C57BL/6N | MGI:3045306 | MGI:6120806 | homozygote | 116 | 0.5520 | 0.5027 | 0.6013 |
| involves: C57BL/6NTac | MGI:3583900 | MGI:5548895 | homozygote | 106 | 0.5552 | 0.5043 | 0.6068 |
| involves: C57BL/6NTac | MGI:1096391 | MGI:5636923 | homozygote | 80 | 0.5575 | 0.5015 | 0.6143 |
| involves: C57BL/6NCrl | MGI:1098802 | MGI:5754588 | heterozygote | 89 | 0.5599 | 0.5020 | 0.6182 |
| involves: C57BL/6NCrl | MGI:1923520 | MGI:5605796 | heterozygote | 127 | 0.5606 | 0.5164 | 0.6058 |
| involves: C57BL/6N | MGI:1891692 | NULL-87B389A49 | homozygote | 58 | 0.5671 | 0.5034 | 0.6338 |
| involves: C57BL/6N | MGI:1920412 | NULL-B804249AA | homozygote | 79 | 0.5671 | 0.5053 | 0.6281 |
| involves: C57BL/6N | MGI:88586 | MGI:5605834 | homozygote | 64 | 0.5712 | 0.5076 | 0.6352 |
| involves: C57BL/6N | MGI:2153041 | MGI:6257715 | homozygote | 52 | 0.5734 | 0.5036 | 0.6491 |
| involves: C57BL/6NCrl | MGI:1923573 | NULL-20A7CB797 | homozygote | 92 | 0.5735 | 0.5122 | 0.6309 |
| involves: C57BL/6N;C57BL/6NTac | MGI:1922656 | MGI:5609354 | heterozygote | 57 | 0.5775 | 0.5082 | 0.6447 |
| involves: C57BL/6N | MGI:1927170 | MGI:5605804 | heterozygote | 57 | 0.5789 | 0.5105 | 0.6461 |
| involves: C57BL/6N | MGI:1935037 | MGI:5568482 | homozygote | 63 | 0.5792 | 0.5148 | 0.6417 |
| involves: C57BL/6N | MGI:2444798 | MGI:5548831 | homozygote | 102 | 0.5799 | 0.5311 | 0.6304 |
| involves: C57BL/6N | MGI:88468 | MGI:5575901 | homozygote | 59 | 0.5812 | 0.5119 | 0.6492 |
| involves: C57BL/6NCrl | MGI:2148202 | MGI:6158484 | homozygote | 73 | 0.5816 | 0.5242 | 0.6408 |
| involves: C57BL/6NCrl | MGI:3613666 | MGI:6158463 | homozygote | 73 | 0.5853 | 0.5310 | 0.6432 |
| involves: C57BL/6N | MGI:2385088 | MGI:5602831 | heterozygote | 59 | 0.5932 | 0.5286 | 0.6570 |
| involves: C57BL/6N | MGI:1922915 | MGI:5603351 | homozygote | 57 | 0.5941 | 0.5215 | 0.6639 |
| involves: C57BL/6N | MGI:1341272 | MGI:5561558 | homozygote | 57 | 0.5994 | 0.5357 | 0.6667 |
| involves: C57BL/6N | MGI:96549 | MGI:5766766 | homozygote | 62 | 0.5995 | 0.5329 | 0.6602 |
| involves: C57BL/6NTac | MGI:1202301 | MGI:5692595 | homozygote | 78 | 0.6014 | 0.5456 | 0.6621 |
| involves: C57BL/6NTac | MGI:104993 | MGI:5692557 | homozygote | 54 | 0.6094 | 0.5386 | 0.6742 |
